# Stimulating microglial neuroinflammation with amyloid-beta oligomers compatible with solid-state NMR structural characterization

**DOI:** 10.64898/2026.10.14.759149

**Authors:** Daniel M Dinakarapandian, Tarunya Rao Sudarshan, Martin N Griffin, Pranay Ramaswamy, Phuong H Nguyen, Tzu-Ying Chiu, Chenxing Li, S. Shirin Kamalaldinezabadi, Jens O Watzlawik, Terrone L Rosenberry, Scott M Stagg, Zhexing Wen, Levi B Wood, Anant K Paravastu

## Abstract

Soluble amyloid-β (Aβ) oligomers are considered the primary neurotoxic species in Alzheimer’s disease, yet the relationship between oligomer structure and biological activity remains poorly understood. Oligomer sample preparations amenable to high-resolution structural characterization have not been tested in cell-based assays, mostly because detergents used to drive oligomer formation are toxic to cells. Here, we describe a detergent-free protocol in which size-exclusion chromatography–isolated Aβ42 monomers spontaneously oligomerize at 37 °C in a concentration- and temperature-dependent manner. Biophysical characterization shows a β-sheet structure by circular dichroism, a self-associating population above 100 kDa by silver-stained SDS-PAGE, and bead-like strings by negative-stain TEM resembling the previously characterized SDS-induced 150 kDa oligomer. Solid-state NMR spectroscopy of selectively ¹³C/¹⁵N-labeled 37 °C oligomers yields well-resolved spectra with β-strand chemical shifts, and two-dimensional ¹³C–¹³C DARR and CHHC experiments provide initial inter-residue distance constraints that reveal both similarities to and differences from the 150 kDa oligomer. Treatment of mouse microglial cells (MMCs) with 50 nM 37 °C oligomers elicited a robust pro-inflammatory cytokine response that exceeded that of monomeric Aβ42, as demonstrated by Luminex multiplex profiling and principal component analysis. hiPSC-derived microglia (iMGs) treated with 1 µM oligomers also separated from vehicle, establishing activity in a human cellular model. Monomeric Aβ42 was inactive in both systems, indicating that inflammatory activity depends on the assembly state of Aβ42 rather than on the presence of peptide alone. These results establish a detergent-free Aβ42 oligomer preparation that is simultaneously compatible with solid-state NMR structural characterization and biologically active in neuroinflammation assays, providing a platform for studying microglial immune responses to oligomers.

## Introduction

We report a detergent-free protocol for producing Aβ42 oligomer samples that are compatible with both solid-state NMR structural characterization and microglial cell assays. Solid-state NMR spectroscopy can provide detailed structural information when samples are sufficient in quantity, homogeneity, and stability. Microglia are the primary innate immune effectors of the central nervous system and likely play a role in the early stages of Alzheimer’s disease (AD)^1,2^. As a result, we are interested in examining cytokine responses to oligomers in the nanomolar to low micromolar range in microglial cell cultures, relevant to human AD pathology. If microglial responses can be measured with well-characterized, structurally defined oligomeric samples, we can investigate structure-pathology relationships in oligomers. However, detergents commonly used to produce structurally defined oligomeric samples can adversely affect cellular responses^3^. To our knowledge, no existing Aβ42 oligomer preparation produced without detergents or organic co-solvents (such as DMSO) has achieved both interpretable solid-state NMR spectra and biological activity at physiologically relevant concentrations.

Aggregation of amyloid-β (Aβ) peptides is a defining feature of AD pathology, yet which specific aggregated species drive neurotoxicity and neuroinflammation remains incompletely understood. It is posited that the accumulation of Aβ peptides derived from proteolytic cleavage of the amyloid precursor protein (APP) by β-and γ-secretases initiates a pathological sequence leading to neuronal death and cognitive decline^4^. Among the Aβ alloforms, Aβ42 is particularly prone to aggregation due to the two additional hydrophobic C-terminal residues (Ile41 and Ala42) absent in the more common Aβ40 isoform. These residues endow Aβ42 with a higher aggregation propensity, forming β-strand structure in the C-terminal region that facilitates intermolecular association. Consistent with this observation, Aβ42 forms oligomeric structures more readily than Aβ40 under comparable conditions^5^. Additionally, there is evidence that implicates soluble oligomeric intermediates, rather than fibrils, as the primary drivers of synaptic dysfunction and neuroinflammation in AD^6–9^. However, oligomer populations can be heterogeneous and present at low abundance, creating a challenge: preparing samples that are simultaneously amenable to high-resolution structural methods and biologically relevant cell-based assays.

Prior studies have established that oligomeric forms of Aβ42, compared to amyloid fibrils, exert distinct effects on microglia, but many questions remain unanswered. The pathological relevance of oligomers is suggested by the greater sensitivity of microglial responses to oligomers than to monomers or fibrils at total Aβ42 concentrations near 50 nM, a range considered physiologically relevant^10,11^. Soluble Aβ oligomers have also been shown to elicit a more potent proinflammatory cytokine response relative to fibrillar Aβ^10^. These studies, however, were conducted on oligomeric preparations lacking structural characterization, making it difficult to determine which structural features of the oligomers govern their biological potency. What remains poorly understood is how specific oligomeric structures, which may differ in receptor binding, β-sheet content, size distributions, and physical properties, relate to inflammatory outcomes. Bridging this gap requires oligomer preparations that can be structurally interrogated while simultaneously serving as stimuli in cell-based assays.

Current methods for producing Aβ42 oligomers suitable for structural study often rely on detergents, such as sodium dodecyl sulfate (SDS)^12,13^ or dodecylphosphorylcholine (DPC)^13,14^. These detergents stabilize assembly pathways by embedding the hydrophobic C-terminal region of Aβ42 within the hydrophobic core of micelles^12,14^. While detergent-assisted protocols have produced the best-characterized oligomers to date, including the SDS-induced 150 kDa oligomer characterized in our work via solid-state NMR^15–17^, the residual detergent in these samples may alter the oligomer surface properties and limit their suitability for cellular experiments. Moreover, the structure of detergent-stabilized oligomers reflects the templating influence of the micelle environment: the antiparallel C-terminal β-sheet (C-sheet; residues 30–42) tetramer formed within SDS or DPC micelles is directly incorporated into the resulting 150 kDa oligomer^13,15,16^. Previous simulation-based attempts to structurally dissect the mechanism underlying the C-terminal antiparallel β-sheet revealed that three glycine residues at positions 33, 37, and 38 create packing voids that are filled by detergent hydrocarbon tails, thereby converting a potentially destabilizing feature into a stabilizing factor^13^. In addition to the C-terminal antiparallel β-sheet, the 150 kDa oligomer also features an N-terminal (residues 11–24) parallel β-sheet, suggesting that coexistence of both types of β-sheets within the same oligomer could be the reason why the oligomer does not grow into fibrils^15^. Cumulatively, structural studies on detergent-assisted oligomers raise questions about whether similar structural motifs exist in oligomers assembled in the absence of detergents. Like the 150 kDa oligomer, it could be possible that detergent-free oligomers may also encompass a parallel and antiparallel β-sheet within the same oligomeric structure. Thus, a detergent-free oligomer preparation, unlike detergent-assisted oligomers, offers a complementary, more physiologically relevant route in which the structure is not induced by an artificial hydrophobic scaffold.

Here, we report the experimental characterization of the “37 °C oligomer.” This contribution includes measurements of the temperature dependence of oligomer formation, the peptide concentration dependence, biophysical properties, and the microglial response. We studied this oligomer in the context of structural characteristics of the 150 kDa oligomer, which we previously produced via a detergent-assisted pathway^15–17^. Size-exclusion chromatography (SEC)–isolated Aβ42 monomers remain stable at 4–6 °C for up to a day but spontaneously form oligomers at 37 °C in a concentration-dependent manner. We show that the 37 °C oligomer has β-strand secondary structure, as confirmed by circular dichroism (CD) spectroscopy, and induces pro-inflammatory cytokine responses in mouse- and human-derived microglial cell cultures. Silver-stained SDS-PAGE reveals a population above ∼100 kDa, and negative-stain TEM reveals structures with morphology like the previously characterized SDS-induced 150 kDa oligomer. These structures, referred to as “strings,” appear to form through the adjacent arrangement of rounded oligomeric units^18^. Solid-state NMR provides initial structural constraints, allowing direct comparison with the detergent-assisted oligomer. By establishing a preparation compatible with both NMR and cellular assays, this work lays the groundwork for correlating oligomer structure with biological activity.

## Results

### WT Aβ42 monomers are stable at 6 °C and form β-sheet-rich oligomers at 37 °C

Fig. 1a summarizes the preparation: SEC-purified monomers are incubated at controlled temperature and the resulting aggregates re-isolated by SEC before downstream analysis.

**Figure 1.**
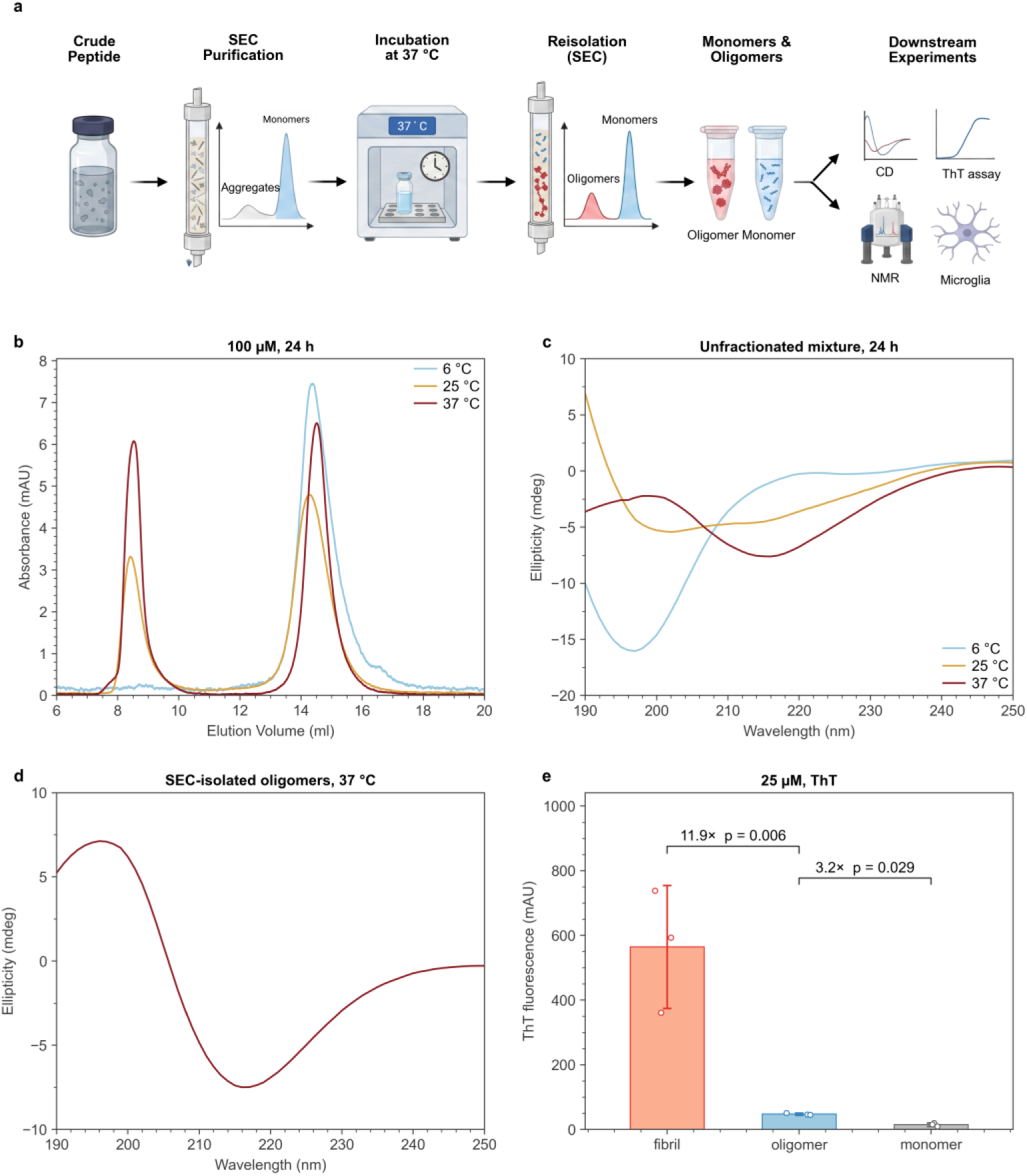
WT Aβ42 forms oligomers between 25 °C and 37 °C. (**a**) Preparation workflow: crude peptide is purified by SEC, the isolated monomer fraction is incubated at 37 °C, and the resulting aggregates are re-isolated by SEC before downstream microglial, NMR, CD and ThT experiments. (**b**) SEC profiles measured for 100 µM SEC-isolated monomers re-injected after 24 h incubation at 6 °C, 25 °C and 37 °C. (**c**) CD spectra of the unfractionated mixture after 24 h incubation at the same three temperatures. (**d**) CD spectrum of SEC-isolated oligomers formed at 37 °C for 24 h. (**e**) ThT fluorescence at 25 µM for three states, each read where that species is defined: fibrils from the monomer arm at its plateau, SEC-isolated oligomers at t = 0, and monomers at t = 0. Bars are the mean of n = 3 with the individual replicates shown; ratios and p values come from Welch t-tests on the log of the replicate values.

We evaluated the stability and propensity for oligomer formation of wild-type (WT) Aβ42 monomers isolated by size-exclusion chromatography (SEC). Synthetic Aβ42 peptide, dissolved at a concentration of 2 mg/mL in 50 mM sodium hydroxide (NaOH), was injected onto a Superdex 75 Increase SEC column run in 20 mM sodium phosphate (NaPi) buffer at pH 7.4. We collected fractions corresponding to the expected monomer elution time from 14.5 to 16.0 mL (Supplementary Fig. 1). We then reinjected those isolated monomers at 100 µM through SEC at different temperatures (Fig. 1b). Circular dichroism (CD) at 50 µM Aβ42 peptide concentration provides further evidence that this elution peak is monomeric; aliquots captured from this fraction exhibited random-coil secondary structure (Fig. 1c), with a dip centered at ∼200 nm wavelength. In contrast, low molecular weight oligomers would be expected to exhibit a secondary structure that is either a β-strand or α-helix. Random coil secondary structure remains stable at 6 °C for 24 h and at 25 °C for 30 min (Fig. 1c). Correspondingly, we observed by SEC only a single monomeric peak with no detectable higher-molecular-weight species with immediate reinjection (time t = 0) or incubation at 6 °C for 24 h. In contrast, WT monomers exposed to 37 °C formed a distinct early-eluting peak at ∼8 mL (Fig. 1b). We assign this early-eluting peak to oligomer formation; early elution corresponds to higher molecular weight. Consistently, an aliquot collected from this early-eluting peak exhibited a CD profile consistent with β-strand secondary structure (Fig. 1d).

Thioflavin-T fluorescence distinguishes the two species^19^: at 25 µM, the SEC-isolated oligomers give a signal 11.9-fold lower than the fibrils formed from monomer at the plateau of the monomer aggregation assay (p = 0.006, Welch t-test on log values, n = 3; Fig. 1e and Supplementary Fig. 2). The oligomer signal is nonetheless 3.2-fold above that of freshly isolated monomer (p = 0.029, same test), placing the oligomer between the monomer and the fibril rather than at either extreme.

### Oligomeric Aβ42 upregulates cytokine expression in mouse and hiPSC-derived microglial cells

Considering that Aβ aggregation state influences microglial activity^1,2,10,20,21^, we next investigated whether oligomeric Aβ42 would elicit distinct cytokine responses compared to monomeric Aβ42. To do so, we applied 50 nM of monomeric Aβ42 and oligomeric Aβ42 to mouse microglial cells (MMC) for 24 h^11^. We then quantified secretion of 15 inflammatory cytokines and chemokines associated with chronic inflammation in the culture medium using a Luminex multiplexed immunoassay (Fig. 2a)^22,23^. Principal component analysis (PCA) identified principal components (PC1 and PC2) that captured the greatest variance in cytokine expression across samples, separating oligomeric Aβ42 from both monomeric Aβ42 and vehicle along PC1 (Fig. 2b, c). PC1 scores were significantly higher for oligomeric Aβ42 than for either monomeric Aβ42 or vehicle (Fig. 2d). The loadings on PC1 reflect the relative contribution of each cytokine to this separation: TNF-α, IP-10, IL-3, and VEGF had the highest absolute loadings (Fig. 2b). Among these top contributors, IP-10 and VEGF were significantly elevated in oligomeric Aβ42 relative to vehicle, and IP-10 and TNF-α were significantly elevated in oligomeric Aβ42 relative to monomeric Aβ42 (Fig. 2e, Supplementary Fig. 3).

**Figure 2.**
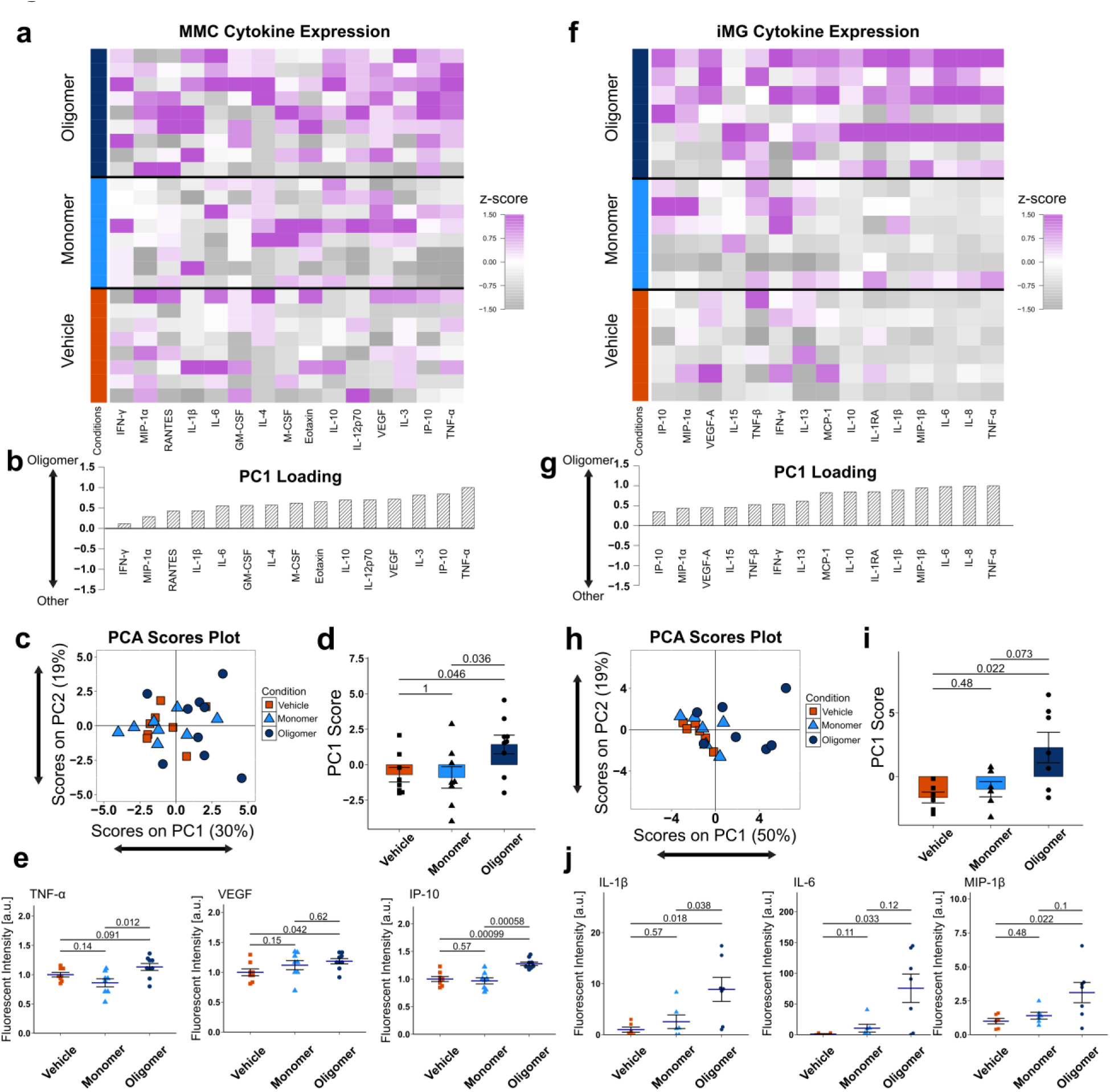
Aβ42 oligomeric solutions induce inflammatory cytokine expression in a mouse microglial cell line and hiPSC-derived microglia. **(a)** Multiplexed Luminex analysis of 15 cytokines (columns) expressed into the MMC culture medium after 24 h of incubation with 50 nM of Aβ monomers, Aβ oligomers, or vehicle (n = 8–9 wells/group). Each row in the z-scored heat map denotes an individual culture sample well. **(b**) PCA identified a variable, PC1, that captured the greatest axis of variation across experimental groups. PC1 consisted of a profile of cytokines that were upregulated with Aβ oligomer. **(c)** Scores plot of principal component analysis (PCA). Each point represents a sample well. PC1 explained 30% of the variance, and PC2 explained an additional 19%. **(d)** Bar plot of PC1 scores for each group (mean ± SEM, Wilcoxon rank sum test). **(e)** Dot plots for individual cytokines (mean ± SEM, Wilcoxon rank sum test). Each dot denotes an individual sample well. All p-values reflect Wilcoxon rank sum tests with Bonferroni adjustment for multiple comparisons. a.u., arbitrary units. **(f)** Multiplexed Luminex analysis of 15 cytokines (columns) expressed into the iMG culture medium after 24 h of incubation with 1 µM of Aβ monomers, Aβ oligomers, or vehicle (n = 6–7 wells/group). Each row in the z-scored heat map denotes an individual culture sample well. **(g)** PCA identified a variable, PC1, that captured the greatest axis of variation across experimental groups. PC1 consisted of a profile of cytokines that were upregulated with Aβ oligomer. **(h)** Scores plot of principal component analysis (PCA). Each point represents a sample well. PC1 explained 50% of the variance, and PC2 explained an additional 19%. **(i**) Bar plot of PC1 scores for each group (mean ± SEM, Wilcoxon rank sum test). **(j)** Dot plots for individual cytokines (mean ± SEM, Wilcoxon rank sum test). Each dot denotes an individual sample well. All p-values reflect Wilcoxon rank sum tests with Bonferroni adjustment for multiple comparisons. a.u., arbitrary units.

To extend these findings to a human cellular model, we differentiated human induced pluripotent stem cells (hiPSCs) into microglia (iMGs) using a previously established protocol^24,25^. We applied 1 µM monomeric Aβ42 or oligomeric Aβ42 to iMGs for 24 h. The higher dose relative to the MMC experiments was chosen because published work in hiPSC-derived microglia demonstrates robust inflammatory and transcriptional responses in the high-nanomolar to low-micromolar range^26–28^. We then quantified the concentrations of 15 inflammatory cytokines and chemokines associated with chronic inflammation secreted into the culture medium using a Luminex multiplexed immunoassay (Fig. 2f)^22,23^. PCA again separated oligomeric Aβ42 from both monomeric Aβ42 and vehicle along PC1 (Fig. 2g, h). PC1 scores were significantly higher for oligomeric Aβ42 than for vehicle, with a comparable but non-significant trend relative to monomeric Aβ42 (Fig. 2i). TNF-α, IL-8, IL-6, and MIP-1β had the highest absolute loadings on PC1 (Fig. 2g). At the level of individual analytes, IL-1β, IL-6, IL-15, and MIP-1β were significantly elevated in oligomeric Aβ42 relative to vehicle, and IL-1β was significantly elevated relative to monomeric Aβ42 (Fig. 2j, Supplementary Fig. 4).

Despite the species difference, the same underlying inflammatory response structure emerged in both the MMCs and iMGs. All analyte loadings were positive on PC1, indicating coordinated upregulation in inflammatory cytokines across the panel. In both systems, oligomer-treated samples separated significantly from vehicle along PC1, with the response to oligomeric Aβ42 exceeding that to monomeric Aβ42.

### Oligomer formation depends on Aβ42 concentration and temperature

The SEC profiles in Fig. 3 demonstrate a nonlinear Aβ42 concentration and temperature dependence for oligomer formation. WT monomers isolated by SEC were incubated at 25 µM, 100 µM, and 250 µM for various durations in 20 mM NaPi buffer, pH 7.4 (Fig. 3a–c). The 250 µM samples showed a stronger oligomeric SEC peak (∼8 mL), consistent with a larger population of higher-molecular-weight aggregates when compared with the 100 µM and 25 µM samples. At 25 µM, aggregation was minimal, although a weak peak consistent with oligomers appeared at longer incubation times. An additional late-eluting SEC peak also appeared at ∼17 mL (Fig. 3c). To assess whether low-abundance oligomers were missed by the initial absorbance measurements at 280 nm, we repeated the 25 µM aggregation assay using UV detection at 210 nm to improve peptide detection sensitivity. These measurements likewise showed little to no detectable oligomer formation (Supplementary Fig. 5). The late-eluting peak was observed in both analyses and was also present following incubation with 0.2% sodium azide (Fig. 3c). LC– MS analysis identified intact Aβ42, arguing against peptide cleavage as the explanation for this peak. The altered elution behavior may reflect a change in peptide conformation or interactions with the column matrix; however, its underlying basis remains unresolved. In addition, incubation at both 37 °C and 25 °C induces aggregation in WT monomers, as shown by the oligomeric peak eluting at ∼8 mL in Fig. 3. However, the increase in the intensity of this oligomeric peak is slower at 25 °C (Fig. 3d) when compared to that at 37 °C (Fig. 3b). Supplementary Fig. 6 facilitates comparison of the areas under the peaks at different injection time points for the SEC profiles observed in Fig. 3. Overall, these data indicate that WT monomer aggregation increases as the incubation temperature and initial monomer concentration increase.

**Figure 3.**
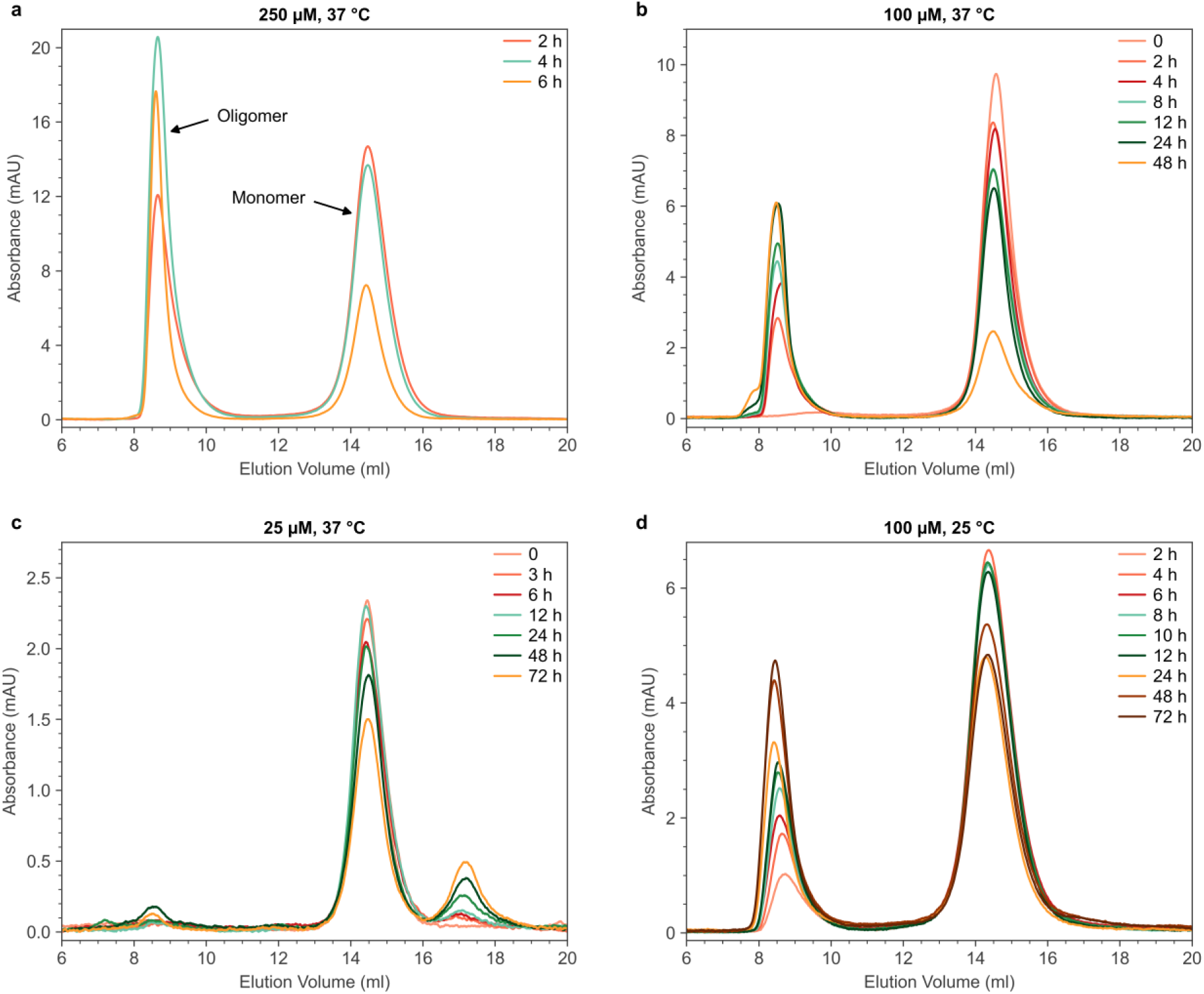
Oligomer formation depends on Aβ42 concentration and temperature. SEC profiles measured for incubation-time-dependent re-injection of WT monomers at (**a**) 250 µM, 37 °C, (**b**) 100 µM, 37 °C, (**c**) 25 µM, 37 °C and (**d**) 100 µM, 25 °C. Absorbance is plotted without normalization so that peak magnitudes remain comparable between time points within each panel.

Temperature-dependent Aβ42 aggregation was further confirmed by CD spectroscopy. At 6 °C, incubated monomers retained their random-coil structure, with a spectral dip below ∼200 nm, even after 24 h of incubation. In contrast, the β-strand signature usually appears at ∼220 nm, and the α-helix signature appears at ∼215 nm. At 25 °C and 24 h incubation time, the sample likely contains a mixture of structures including molecules in random coil and β-strand conformations. At 37 °C and after 24 h of incubation, the monomers converted almost entirely to a β-strand signature (Fig. 1c). CD spectra of SEC-isolated oligomers obtained from monomers incubated for 24 h at 37 °C further confirmed a well-defined β-strand minimum at ∼220 nm (Fig. 1d). We subsequently refer to the higher-molecular-weight aggregates that elute at ∼8 mL in our SEC figures as “37 °C oligomers.” Cumulatively, SEC and CD results demonstrate that WT monomers are initially stable after purification but exhibit increased aggregation at elevated temperature and monomer concentration.

### 37 °C oligomers exhibit molecular weight bands above 100 kDa in SDS-PAGE

To qualitatively assess the molecular weight distribution of the SEC-purified fractions of the Aβ oligomers collected in the range of elution volume from 8–10 mL, we performed silver-stained SDS-PAGE analysis. First, we reconstituted three lyophilized oligomer fractions in PBS: the SDS-induced 150 kDa fraction (50 µL), the S1 fraction (20 µL), and the S2 fraction (50 µL). The fractions S1 and S2 represent independent batches of temperature-induced 37 °C oligomers.

We loaded 1 µL and 2 µL of each sample without prior concentration determination, and electrophoresis was performed at room temperature (25 °C).

Silver staining revealed distinct differences between the temperature-induced and SDS-induced oligomers (Fig. 4a). The S2 batch showed the strongest bands, with signal extending to molecular weights above 100 kDa, which exceeded the molecular weights exhibited by 150 kDa oligomers (the assignment of 150 kDa as the molecular weight of this oligomer was made previously by light scattering^29^). The S1 batch displayed similar banding patterns but with lower intensity, consistent with a lower peptide concentration. Both detergent-free oligomer batches formed larger assemblies than the SDS-induced 150 kDa fraction, which showed weaker staining with bands concentrated near 150 kDa. Band intensities scaled proportionally with loading volume (1 vs. 2 µL), confirming that differences between samples reflect peptide concentration rather than staining artifacts. All three preparations contained multiple oligomeric species, but the detergent-free oligomers showed a clear shift toward higher molecular weights compared to the SDS-induced sample. Because SDS-PAGE is performed under denaturing conditions, the higher molecular weight bands observed for 37 °C oligomers may reflect either intrinsically larger assemblies or greater resistance to SDS-induced dissociation relative to the 150 kDa oligomer. A tetrameric (∼18 kDa) band was consistently observed across all samples; however, this species may result from dissociation of larger assemblies by residual SDS during electrophoresis rather than representing a native oligomeric population.

**Figure 4.**
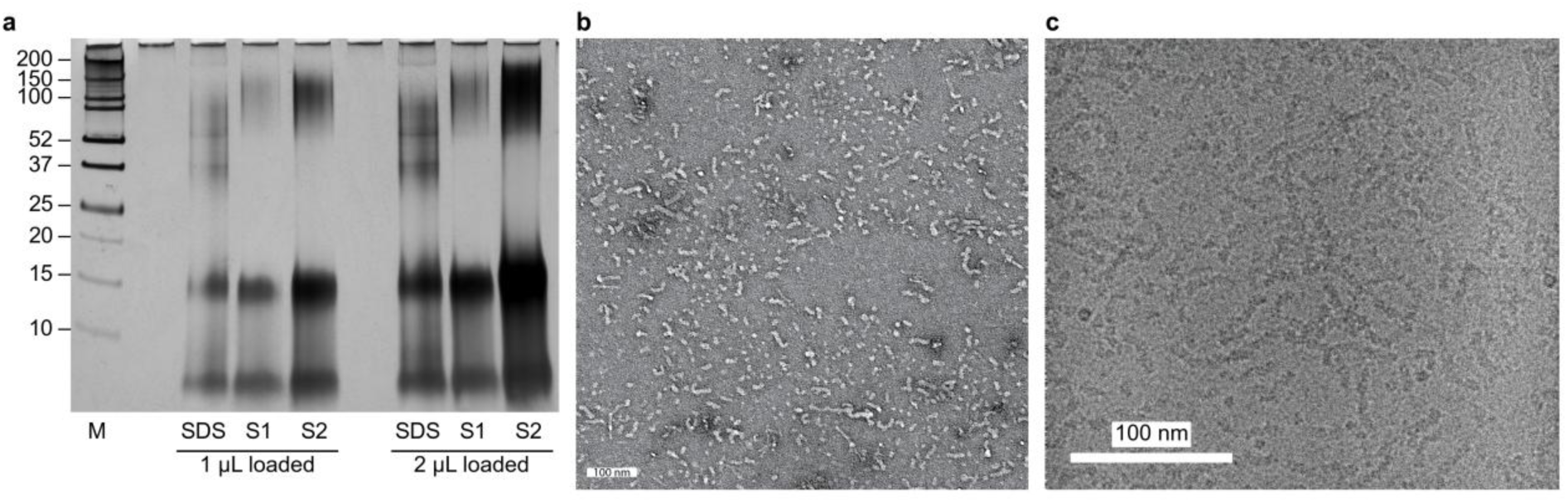
Size distribution and morphology of 37 °C oligomers. (**a**) Silver-stained SDS-PAGE of SEC-fractionated Aβ42 oligomers, run at room temperature (25 °C). Lane SDS is the SDS-induced 150 kDa fraction; lanes S1 and S2 are independently prepared 37 °C oligomers, each loaded at 1 µL and 2 µL without prior concentration determination. Lane M is the Bio-Rad protein marker and unmarked lanes are empty. All sample lanes (SDS, S1 and S2) were lyophilized and reconstituted in PBS before electrophoresis. (**b**) Negative-stain TEM image showing rounded units arranged in strings. (**c**) Cryo-EM micrograph of 37 °C Aβ42 oligomers in string form. Oligomers were prepared from a 250 µM Aβ42 monomer solution incubated at 37 °C for 4 h, isolated by size-exclusion chromatography (SEC), lyophilized following removal of approximately 70% of the 20 mM NaPi buffer, and resuspended in 10 mM NaPi prior to vitrification and cryo-EM imaging on a Titan Krios at 59,000× magnification. Scale bar, 100 nm.

### Cryo-EM shows that 37 °C oligomers exhibit similar morphologies to the SDS induced-150 kDa oligomers: Circular particles and ‘string-like’ aggregates

We performed cryo-EM imaging (Fig. 4b, c) to characterize the sizes and shapes of 37 °C oligomers, and check for the presence of fibrils. The sample used for cryo-EM was prepared from an SEC-isolated 250 µM Aβ42 monomer solution incubated at 37 °C for 4 h, isolated by a second SEC injection (oligomer peak at ∼8 mL), and lyophilized after removal of ∼70% of the NaPi buffer (details in Materials and Methods). We observed similar nanoscale morphologies to what we reported previously for the 150 kDa oligomer: no amyloid fibrils, circular particles with diameters of 6–7 nm, and elongated structures resembling “strings.”^18^ We previously introduced the term strings to describe elongated structures that likely differ from amyloid fibrils. We also avoid the term “protofibrils” because we observed no evidence that strings are on pathway to fibril formation. Instead, we propose that strings are the result of self-assembly of oligomers. While oligomers appear spherical in EM images, their assembly into strings would indicate anisotropic structures.

### Solid-state NMR spectroscopy indicates an interpretable structure with similarities and differences compared to the previously investigated 150 kDa oligomer

Our goal with NMR measurements on the 37 °C oligomer samples was to evaluate whether the oligomer is ordered enough for structural analysis and whether it carries any of the structural motifs previously identified in the 150 kDa oligomer. To facilitate a structural comparison between the two oligomers, we uniformly labeled six residues L17, F20, A30, M35, V36, and I41 in the Aβ42 sequence with ¹³C and ¹⁵N material and prepared 37 °C oligomers (see Materials and Methods). The labels were strategically chosen to identify if the 37 °C oligomer carried the same underlying structural arrangement as the 150 kDa oligomer. The 150 kDa oligomer has been previously structurally characterized through solid-state NMR and cryo-EM measurements^15,16,18^ and contains a combination of antiparallel and parallel β-sheets^15,16^ (see Fig. 5a), which sets it apart from the widely characterized parallel β-sheet fibrils. Fig. 5a shows a cartoon representation of the 150 kDa oligomer along with the exact residues in the C-terminal and N-terminal regions of the Aβ42 peptide that participate in the coexisting parallel and antiparallel β-strands. By placing ¹³C labels at L17 and F20, we can check if the 37 °C oligomers also form the n→n+3 parallel β-strand arrangement found in the N-terminal region of the 150 kDa oligomer. ¹³C labels at A30 and I41 allow us to probe the C-terminal antiparallel strand in the 150 kDa structure. Lastly, labels at M35 and V36 help us assess if the C-terminal antiparallel β-strand would stack against the N-terminal parallel β-strand, which would be reflected by proximity with L17 as observed in the 150 kDa structure. Fig. 5b, c encompasses these anticipated proximities 2D NMR could detect if the 37 °C oligomer was structurally similar to the 150 kDa oligomer. Thus, our labeling strategy allows for direct comparison between the 37 °C oligomers and the previously characterized 150 kDa oligomer.

**Figure 5.**
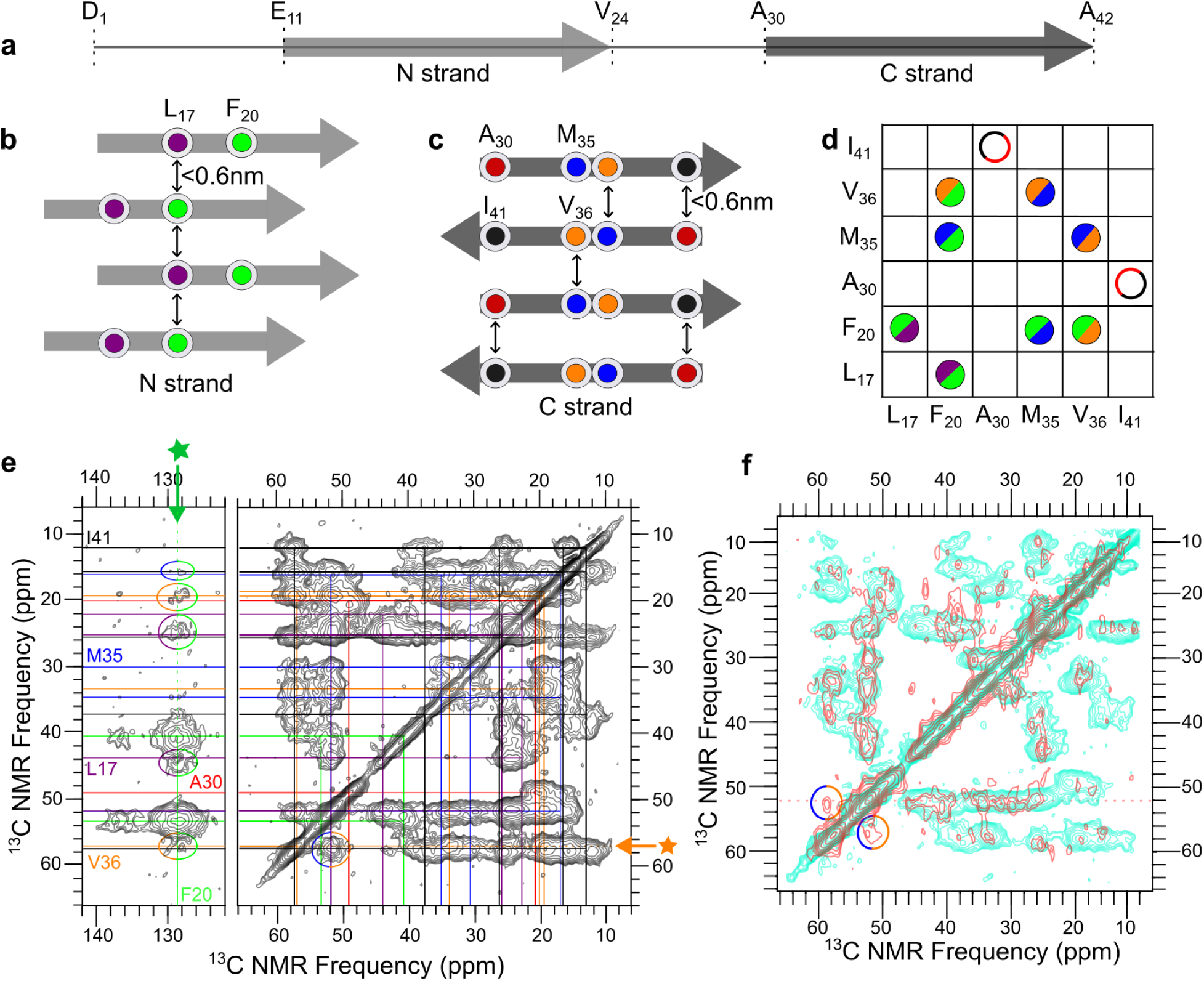
Solid-state NMR constraints on the 37 °C oligomer. (**a**) Aβ42 sequence with the N-terminal and C-terminal β-strand regions assigned in the 150 kDa oligomer. (**b**) N-terminal parallel β-strand arrangement, with L17 (purple) and F20 (green) marked; arrows indicate separations below 0.6 nm at which 2D NMR proximities may be detected. (**c**) C-terminal antiparallel β-strand arrangement, with A30 (red), M35 (blue), V36 (orange) and I41 (black) marked. (**d**) Proximities detected in the 37 °C oligomer. Bi-colored semicircles denote detected crosspeaks; open red circles denote expected proximities that were not detected. (**e**) 2D ¹³C–¹³C DARR spectrum at 500 ms mixing time, with assignments drawn as colored lines and detected crosspeaks circled. Stars mark the frequencies of the slices in Supplementary Fig. 8. (**f**) Overlay of the 2D CHHC spectrum (red) on the 20 ms 2D DARR spectrum (cyan); circles mark the M35 Cα–V36 Cα proximity.

Repeatability of the preparation was checked by ¹H–¹³C CPMAS on two independently prepared oligomer samples, whose spectra overlap (Supplementary Fig. 7).

To define residue-specific NMR chemical shifts and evaluate structural similarity between the 150 kDa oligomer and 37 °C oligomer, we conducted 2D ¹³C–¹³C DARR measurements on the 37 °C oligomer at a shorter mixing time^30^. Short-mixing DARR experiments primarily detect magnetization transfer between ¹³C nuclei within the same residue, allowing us to determine accurate chemical shifts for each labeled carbon site. Using a 2D DARR experiment at a mixing time of 20 ms (see Supplementary Fig. 8), we obtained exact chemical shift assignments for carbon atoms in L17, F20, A30, M35, V36 and I41 indicated using purple, green, red, blue, orange and black solid lines respectively. These assignments enabled us to compute a secondary shift analysis^31^ for the 37 °C oligomer. The corresponding chemical shifts, full-width-at-half-maximum linewidths, and secondary shifts (Δδ) are reported in Table 1. For each labeled residue, the CO, Cα, and Cβ secondary shifts follow the expected –, –, + pattern associated with β-strand structure. This consistent pattern across all labeled residues confirms that the 37 °C oligomer adopts a β-strand–rich configuration, like the 150 kDa oligomer. Moreover, the linewidths of the 37 °C oligomer more closely resemble those reported for the 150 kDa oligomer than those reported for Aβ42 fibrils, suggesting that both oligomers exhibit a comparable level of structural order. However, similar linewidths alone do not correspond to identical structures on a molecular level. To get further insight, we calculated an RMSD (root mean square deviation for corresponding ¹³C sites) between the two sets of chemical shifts, using the 27 carbon sites assigned to the same atom in both oligomers (37 °C shifts from Table 1; 150 kDa shifts from Supplementary Table S1 of Gao et al.^15^), and obtained a value of 0.39 ppm. A value of this size is consistent with similar local conformations at the labeled residues. For comparison, the same calculation against two Aβ42 fibril structures, Colvin et al.^32^ (PDB 5KK3, BMRB 30121) and Gremer et al.^33^ (PDB 5OQV, BMRB 27212), gives RMSDs of 1.1–1.2 ppm over the C′, Cα and Cβ sites after removal of any uniform referencing offset, more than 2.5 times the 0.43 ppm obtained against the 150 kDa oligomer over the same 18 backbone sites. The two fibril structures differ from each other by 1.4 ppm when calculated in the same way.

**Table 1.** Secondary Shift Analysis for the 37 °C oligomer. Each entry corresponds to the NMR chemical shift (ppm) of a uniformly ¹³C-labeled carbon atom; the full width at half maximum linewidths are in parentheses. Δδ values depict secondary chemical shifts (ppm) for CO, Cα, and Cβ carbon atoms. The F20 Cγ position overlaps the expected spinning-sideband position and is therefore left blank; the F20 aromatic-ring signal (128.7 ppm) is listed under Cδ.

| Residue | CO | Cα | Cβ | Cγ (1) | Cγ (2) | Cδ |
| --- | --- | --- | --- | --- | --- | --- |
| L17 | 172.4 (2.8) | 52 (2.8) | 44 (2.8) | 25.3 (2.3) | 22.2 (2.8) |  |
| Δδ | −3.5 | −1.4 | 3.3 |  |  |  |
| F20 | 172.2 (1.9) | 53.6 (2.9) | 40.7 (4.7) |  |  | 128.7 (4.3) |
| Δδ | −1.9 | −2.4 | 2.8 |  |  |  |
| A30 | 173.4 (3.3) | 49.2 (3.8) | 20.1 (5.4) |  |  |  |
| Δδ | −2.7 | −1.6 | 2.7 |  |  |  |
| M35 | 171.9 (3) | 52.2 (3) | 34.8 (3.3) | 30.2 (2.7) | 16.2 (2.3) |  |
| Δδ | −2.7 | −1.6 | 3.6 |  |  |  |
| V36 | 172 (3.2) | 57.3 (3.8) | 33.5 (2.9) | 19.4 (2.6) | 18.7 (0.7) |  |
| Δδ | −2.6 | −3.2 | 2.3 |  |  |  |
| I41 | 172 (3.2) | 57.7 (2.9) | 37.4 (5.8) | 25.7 (2.9) | 15.8 (2) | 12.2 (2.4) |
| Δδ | −2.7 | −1.7 | 0.3 |  |  |  |

To identify site-specific intermolecular proximities within the 37 °C oligomer, we performed ¹³C–¹³C DARR experiments using a 500 ms mixing time. Long-mixing DARR enables magnetization transfer between ¹³C nuclei within roughly 0.6 nm, allowing us to detect structural proximities between labeled residues. In contrast, short-mixing DARR (see Supplementary Fig. 8) primarily reports on intra-residue or intramolecular contacts. Throughout this work, we refer to intermolecular proximities as “crosspeaks.” Fig. 5e shows the long-mixing DARR spectrum with bi-colored circles marking the detected crosspeaks. First, we can see an L17–F20 crosspeak in Fig. 5e, highlighted in purple-green circles. A comparison between horizontal slices taken at the F20 Cδ position from the two DARR experiments showed a peak at 25.3 ppm corresponding to the L17 Cγ signal only in the long mixing spectrum (see Supplementary Fig. 8c). These results confirm the presence of an intermolecular proximity between L17 and F20, which is characteristic of the n→n+3 parallel β-strand arrangement in the N-terminal region of the 150 kDa oligomer. Second, we did not detect an A30–I41 signal, which is one of the defining C-terminal antiparallel contacts in the 150 kDa oligomer. Although we observed M35–V36 crosspeaks (denoted by blue-orange circles in Fig. 5e), these residues are adjacent in sequence, so this proximity alone does not define the underlying strand arrangement.

In the 150 kDa oligomer, labeled residues A30 and I41 define the edges of the C-terminal antiparallel β-sheet, with the sheet centered around V36. It is possible, however, that an antiparallel arrangement could still exist in the 37 °C oligomer but involve only the central portion of the strand, with the edge residues no longer participating. To test this possibility, we focused on M35 and V36, which lie near the center of the C-terminal strand and could reveal an antiparallel arrangement even if the terminal residues are absent from the interaction. Unlike DARR, CHHC relies on proton-driven Cα–Cα magnetization transfer and is suited for identifying the short Cα–Cα distances characteristic of antiparallel β-strands^34^. 2D CHHC revealed a distinct M35–V36 crosspeak (see Fig. 5f), shown by the orange and blue circles. A horizontal slice taken at the M35 Cα position revealed a signal at approximately 58 ppm corresponding to the V36 Cα, and this peak was not present in the short-mixing DARR spectrum (see Supplementary Fig. 9b). This confirms that the residues M35 and V36 are arranged antiparallel to one another. Importantly, this was the only antiparallel contact we detected; neither A30 nor I41 showed CHHC crosspeaks. In the 150 kDa oligomer, the C-terminal antiparallel β-strand spans from A30 to I41, but our data indicate that the antiparallel component in the 37 °C oligomer is more limited and does not extend to these terminal residues. Third, we observed intermolecular crosspeaks such as M35–L17 and V36–L17 in Fig. 5e and Supplementary Fig. 8, which likely arise from stacking between the N-terminal parallel strand and the shorter C-terminal antiparallel strand, similar to what has been reported for the 150 kDa species. Fig. 5d illustrates the contacts we observed from our 2D DARR and 2D CHHC experiments. Overall, the results suggest that the 37 °C oligomer and the 150 kDa oligomer may share broad structural features, like the presence of both parallel and antiparallel β-strand elements. However, our 2D NMR results on the 37 °C oligomer indicate that its β-strand boundaries are not identical to those of the 150 kDa oligomer, and additional experiments will be needed to fully determine how the structures differ.

## Discussion

The central objective of this work was to develop a detergent-free preparation of Aβ42 oligomers compatible with both solid-state NMR structural characterization and cellular assays. We demonstrate that temperature-controlled incubation of SEC-purified monomers at 37 °C achieves this goal. The resulting 37 °C oligomers yield interpretable two-dimensional ¹³C–¹³C DARR and CHHC spectra with linewidths comparable to the SDS-induced 150 kDa oligomer, confirming sufficient structural order for solid-state NMR analysis without detergent stabilization. The same preparation elicits a robust inflammatory cytokine response in MMCs at 50 nM, significantly exceeding the response to monomeric Aβ42 across multiple cytokines, and drives a similar response in iMGs at 1 µM, with TNF-α among the leading contributors in both systems. This dual compatibility distinguishes the protocol from detergent-assisted preparations, which may be unsuitable for cell studies, and from structurally uncharacterized oligomeric preparations commonly used in biological assays.

There is a marked temperature dependence of oligomerization: monomers are stable at 6 °C, partially convert at 25 °C, and undergo near-complete conversion to β-sheet-rich assemblies at 37 °C. Two explanations may be considered: a kinetic effect in which temperature provides energy to cross a nucleation barrier, or a thermodynamic effect rooted in the hydrophobic effect. The kinetic explanation is unlikely to be based solely on β-sheet propagation, because amyloid fibril seeds propagate readily at room temperature^35–37^. This phenomenon indicates that the barrier to templated β-sheet extension is accessible at lower temperatures, at least for in-register parallel β-sheets common for fibrillar structures^35,36^. Furthermore, monomers are already in a random-coil conformation at 6 °C, with no folded structure to unfold. The temperature dependence seems more indicative of the hydrophobic effect, which strengthens with increasing temperature as the entropic cost of maintaining ordered hydration shells around nonpolar surfaces rises^37,38^. The C-terminal region of Aβ42, carrying residues A30–A42, is substantially more hydrophobic than the N-terminal region and contributes the dominant thermodynamic driving force for aggregation^39,40^. Whether the 37 °C oligomers represent thermodynamically stable species or kinetically trapped assemblies, and how incubation temperature affects internal structure, remain open questions.

Solid-state NMR reveals both structural similarities and differences between the 37 °C oligomer and the SDS-induced 150 kDa oligomer. The 500 ms DARR spectrum contains an L17–F20 crosspeak that is consistent with the n→n+3 registry-shifted parallel β-strand arrangement in the N-terminal region, the same motif identified in the 150 kDa oligomer and distinct from the in-register parallel β-sheets in all characterized Aβ42 fibril structures^15,32,33,41^. The conservation of this registry across two entirely independent preparation methods may reflect an intrinsic structural preference of the Aβ42 sequence rather than an artifact of detergent templating. This observation is consistent with proposals that out-of-register β-sheet organization may distinguish oligomeric from fibrillar pathways^42,43^. The molecular interactions that favor this specific registry offset remain under investigation. In contrast, the C-terminal antiparallel arrangement differs: CHHC detects an M35–V36 Cα–Cα proximity but the A30–I41 contact that spans the full antiparallel C-sheet in the 150 kDa oligomer is absent. This result is consistent with the glycine-hole mechanism described for the detergent-stabilized oligomer: three glycine residues at positions 33, 37, and 38 create packing voids within the C-terminal antiparallel β-sheet that are filled by detergent hydrocarbon tails in the 150 kDa oligomer, stabilizing the full A30–I41 span^13^. Without detergent, these voids remain unfilled, and the edge residues A30 and I41 lose their stabilizing contacts, while the central M35–V36 antiparallel arrangement survives because the flanking residues at these positions have bulkier side chains that provide sufficient packing in the absence of a hydrophobic scaffold. The chemical shift RMSD of 0.39 ppm supports shared architecture but non-identical atomic-level structure. Against the two fibril structures the RMSDs are more than twice as large, so at the labeled residues the 37 °C oligomer is locally closer to the 150 kDa oligomer than to the fibrils.

The 37 °C oligomers are biologically active in both mouse and human microglia, establishing that our preparation protocols are compatible with cellular assays. In both MMCs and iMGs, oligomer-treated samples separated from vehicle along PC1, with all analytes loading positively. In MMCs, IP-10 and VEGF were elevated relative to vehicle and IP-10 and TNF-α relative to monomer. In iMGs, IL-1β, IL-6, IL-15, and MIP-1β were elevated relative to vehicle and IL-1β relative to monomer. Monomeric Aβ42 produced no detectable cytokine response relative to vehicle in either system, at the level of PC1 or any individual analyte. The idea that oligomers are more potent inflammatory stimuli than monomers is well supported in the literature^10,11,44^, and our MMC data showed a significant difference in scores on PC1 between monomeric and oligomeric Aβ42. In iMGs, the direct oligomer versus monomer contrast on PC1 did not reach significance, but oligomer treatment separated from vehicle while monomer treatment did not, indicating a difference in inflammatory signaling between vehicle and oligomer that was absent between vehicle and monomer. In both cases, oligomeric Aβ42 produced an elevated inflammatory signature relative to vehicle, consistent with our prior work linking Aβ42 aggregation propensity to microglial cytokine responses^20,21^. Taken together, these results indicate that inflammatory activity depends on the assembly state of Aβ42 rather than on the presence of Aβ42 peptide alone.

The two systems differed in which analytes drove the oligomer-specific response, with MMCs weighted toward IP-10 and VEGF and iMGs toward an IL-1β/IL-6/MIP-1β axis. The iMG experiments were designed to establish whether the preparation elicits an inflammatory response in human microglia, and the dose was selected accordingly. We used 1 µM, below a 3 µM concentration at which conventionally prepared Aβ oligomers have been shown to activate hiPSC-derived microglia over a comparable 24 h exposure^26,27^. Because the two systems differ in both cell origin and dose, the source of the divergence in cytokines cannot be resolved here. Species-specific differences in Aβ engagement are one possible contributor, but dose cannot be excluded. Given that both cell types received the same oligomer preparation, the divergence observed here cannot reflect differences among conformers. Whether structurally distinct oligomers elicit correspondingly distinct inflammatory programs is a separate question, one requiring cell background to be held fixed while the conformer is varied, as in our prior work on the Aβ N27D and Q15E variants^20,21^. If they do, the neuroinflammatory environment in the AD brain may shift as the balance of aggregation species changes over the course of disease^1^.

The present solid-state NMR results support the interpretation that the 37 °C oligomer, like the 150 kDa oligomer, is characterized by co-existence of parallel and antiparallel β-sheets. Although additional structural measurements are required to determine the precise β-strand stretches within the 37 °C oligomer, the present evidence is consistent with our previous proposal that oligomer sizes are limited by geometric frustration associated with β-strand organization^15,42,45–48^. We further suggest that the n→n+3 registry-shifted parallel β-strand motif in the N-terminal region may be intrinsic to the Aβ42 sequence rather than an artifact of detergent templating. Finally, both 37 °C oligomers and 150 kDa oligomers have antiparallel arrangements of C-terminal strands, though the C-terminal strand appears shorter in the 37 °C oligomer. While the β-strand organization in the two oligomers appears similar, they do not share the same structure, indicating that multiple oligomerization pathways can yield biologically distinct species. To our knowledge, this is the first Aβ42 oligomer preparation produced entirely without detergents or organic co-solvents following isolation of monomers that yields interpretable two-dimensional solid-state NMR spectra and elicits a robust inflammatory response at physiologically relevant concentrations. Future studies can detail microglial and neuronal responses to the 37 °C oligomer, and more detailed solid-state NMR measurements will further define the molecular structure. While detergents preclude cellular-mechanistic comparison with the 150 kDa oligomer, biophysical and structural comparisons remain possible.

## Materials and Methods

Full details of the materials and methods are provided in the Supplementary Information.

## Acknowledgements

The authors acknowledge the use of instruments at the NMR center at the Georgia Institute of Technology. We also thank Dr. Dexter Dean (School of Chemistry and Biochemistry, Georgia Institute of Technology), Professor Vijay Rangachari (School of Mathematics and Natural Sciences, University of Southern Mississippi), and Dr. Jhinuk Saha (Institute of Molecular Biophysics, Florida State University) for fruitful discussions.

## Author Contributions

A.K.P. and L.B.W. supervised the research. D.M.D., T.R.S., M.N.G., L.B.W., and A.K.P. designed the experiments. D.M.D., T.R.S., M.N.G., P.R., T.Y.C., C.L., P.H.N., J.O.W., and S.S.K. conducted experiments; D.M.D., T.R.S., M.N.G., C.L., P.H.N., and S.S.K. analyzed data. All authors discussed the results and contributed to the final manuscript. D.M.D., T.R.S., M.N.G., T.L.R., S.M.S., Z.W., L.B.W. and A.K.P. helped revise and finalize edits to the manuscript.

