## Supplemental material for "Stimulating microglial neuroinflammation with amyloid-beta oligomers compatible with solid-state NMR structural characterization"

### Supplementary Information

### Materials and Methods

Reference numbers in the Supplementary Information refer to the reference list of the main text.

#### Materials and peptide synthesis

Aβ42 peptides, unlabeled or selectively ¹³C/¹⁵N-labeled, were synthesized by Fmoc solid-phase peptide synthesis on a CEM Liberty Blue 2.0 synthesizer (CEM, Matthews, NC) using isotope-labeled amino acids from Cambridge Isotope Laboratories (Tewksbury, MA), cleaved with trifluoroacetic acid, verified and purified, as described previously^15,17,20^.

#### Monomer Isolation Using SEC and Subsequent Incubations

Following established protocols^17,49,50^, 6 mg of synthetic Aβ42 peptide was dissolved in 3 mL of 50 mM NaOH and allowed to sit at room temperature for 20 min. The solution was then injected onto a Superdex 75 Increase 10/300 column (Cytiva) equilibrated with 20 mM sodium phosphate buffer (pH 7.4) and operated at a flow rate of 0.8 mL/min. Aβ42 monomers eluted at approximately 15 mL, corresponding to the expected 4.5 kDa species on the SEC profile (Supplementary Fig. 1), and the 14.5–16.0 mL fractions were collected for further experiments. Monomer concentrations were determined using 280 nm absorbance. The collected monomer fractions were then diluted to the desired concentrations using 20 mM sodium phosphate buffer.

To assess the effect of temperature and concentration on oligomer formation, SEC-isolated Aβ42 monomers were incubated at the target temperature for defined time intervals. Following incubation, the samples were reinjected onto the same SEC column to evaluate the presence and intensity of any oligomeric species that formed during the incubation period.

#### Aβ42 Preparation for Microglial Cellular Response

Aβ42 monomers isolated from the SEC were diluted to a concentration of 250 µM and incubated at 37 °C for 4 h and subsequently injected into the SEC column. The oligomers formed, which elute at ~8 mL volume, were collected. These oligomers were kept on ice and diluted with cold 20 mM sodium phosphate buffer to a final concentration of 50 nM for the mouse microglial cytokine response study and 1 µM for the hiPSC-derived microglial cytokine response study.

#### Microglial Cell Line

The MMCs used in this study were generated by immortalizing primary C57BL/6 microglia using the J2 retrovirus^51^.

#### Cell Conditioning and Media Collection for Mouse Microglia

Mouse microglial cells (MMCs) were cultured in DMEM-F12 medium supplemented with 10% fetal bovine serum and 1% Penicillin-Streptomycin until they reached 70% confluency. Cells were stored in an incubator with 95% air and 5% CO_2_ atmosphere at 37 °C. Upon reaching 70% confluency, cells were seeded into 12-well plates and incubated for 24 h. Cells were treated with either 50 nM Aβ42 monomers, 50 nM Aβ42 oligomers, or vehicle (20 mM NaPi) for 24 h followed by medium collection. Collected medium was centrifuged at 13,000 rpm for 10 min at 4 °C, then the supernatant was collected and stored at −80 °C. Multiplexed cytokine quantification was conducted using the Milliplex® MAP Mouse Cytokine/Chemokine 16-Plex kit (Eotaxin, GM-CSF, IFN-γ, IL-1β, IL-2*, IL-3, IL-4, IL-6, IL-10, IL-12p70, IP-10, M-CSF, MIP-1α, RANTES, TNF-α, and VEGF) (Millipore Sigma MCYTOMAG-70K). Cytokines marked with an asterisk did not fall within a linear range of bead fluorescent intensity vs. protein concentration and were not included in the analysis.

#### iPSC Source and Maintenance

The WTC-11 human iPSC line was obtained from the Coriell Institute for Medical Research (GM25256; healthy 30-year-old male donor). iPSCs were maintained on Matrigel-coated plates in mTeSR1 with daily media changes and weekly passaging.

#### iPSC Microglia Differentiation

Hematopoietic progenitor cells (HPCs) were generated from iPSCs using the STEMdiff Hematopoietic Kit according to the manufacturer’s instructions. HPCs were collected on days 12 and 14 of differentiation and plated at 100,000 cells per well on Matrigel-coated 6-well plates. Microglial differentiation was carried out as previously described^24,25^. From day 12 through day 37, cells were cultured in microglial differentiation medium consisting of DMEM/F12 supplemented with 2% B27, 2% ITS-G, 0.5% N2, 1% GlutaMAX, 1% non-essential amino acids, 0.05% insulin, 0.0018% monothioglycerol, 100 ng/mL recombinant human IL-34, 50 ng/mL recombinant human TGF-β1, and 25 ng/mL recombinant human M-CSF. Fresh medium was added at 1 mL per well every other day, with supernatant volume reduced by centrifugation twice over the differentiation period. On day 37, cells were harvested with Accutase and replated at 8,000 cells per well in 96-well plates. From day 37 onward, maturation medium consisted of the differentiation medium above supplemented with 100 ng/mL recombinant human CX3CL1 and 100 ng/mL recombinant human CD200.

#### Cell Conditioning and Media Collection for iPSC Microglia

On day 40, cells were treated with either 1 µM Aβ42 monomers, 1 µM Aβ42 oligomers, or vehicle (20 mM NaPi) for 24 h followed by medium collection. Collected medium was centrifuged at 13,000 rpm for 10 min at 4 °C, then the supernatant was collected and stored at −80 °C. Multiplexed cytokine quantification was conducted using a custom 16-plex Milliplex® Human Cytokine/Chemokine/Growth Factor Panel A kit (IFN-γ, IL-1β, IL-1RA, IL-2*, IL-6, IL-8, IL-10, IL-13, IL-15, IP-10, MCP-1, MIP-1α, MIP-1β, TNF-α, TNF-β, and VEGF-A) (Millipore Sigma HCYTA-60K-16C). Cytokines marked with an asterisk did not fall within a linear range of bead fluorescent intensity vs. protein concentration and were not included in the analysis.

#### Multivariate Analysis of Cytokines

Data were analyzed and figures were generated via RStudio (Boston, MA) using the R programming language. The tidyverse collection of packages was used to conduct data processing. R package heatmap3 was used to generate heatmaps; R packages ggplot2, ggpubr, ggbeeswarm, and ggprism were used to generate dotplots and barplots. Clustering was conducted using the hclust function of the stats package in R using Euclidean distance in the unweighted pair group method with arithmetic mean. For protein analysis, Wilcoxon rank-sum tests were performed, and Bonferroni adjustment was applied to correct for multiple comparisons.

Multivariate cytokine data were analyzed using Principal Component Analysis (PCA) to identify axes, called principal components (PCs), that capture the greatest variance in cytokine expression across samples. PCA was performed using the R package ropls^52^, with all data z-scored prior to analysis. Two principal components were extracted, and 5-fold cross-validation was applied to evaluate model robustness. PCA loadings were used to represent the contribution of each analyte to the observed variance.

#### Circular Dichroism spectroscopy

Aβ42 monomers isolated from SEC were diluted to 50 µM using cold MQ-grade water and kept on ice until CD measurements were performed. CD spectra were collected at 6 °C using a Chirascan™-plus spectrometer (Applied Photophysics, Ltd.) after baseline correction with MQ water lacking peptide. Monomeric state was first confirmed by measuring CD at 6 °C, followed by CD measurements at the required experimental temperatures. All measurements were performed in quartz cuvettes with a 0.1 mm path length with the temperature of the measurements set as per requirements. Each spectrum represents the average of three repeats, which were acquired to confirm measurement repeatability.

#### Thioflavin-T fluorescence assay

Thioflavin-T (ThT) fluorescence kinetics were acquired on a BioTek Synergy H1 in 96-well, black-walled, clear-bottom microplates. Freshly SEC-isolated Aβ42 monomers and oligomers were diluted in 20 mM NaPi buffer, pH 7.4, to a final peptide concentration of 25 µM and added to wells containing ThT to give a final ThT concentration of 20 µM in the well. Plates were sealed with optical-clear film and incubated at 37 °C with shaking, and readings were taken at 15 min intervals. Three replicate wells were measured per sample in each independent experiment, and the per-well kinetic traces were buffer-subtracted and averaged within samples prior to plotting. The fibril value in Fig. 1e is the mean over the plateau (the last 10 h of the run) of the monomer samples, which aggregate under these conditions (Supplementary Fig. 2).

#### Lyophilized oligomer preparation protocol

Aβ42 peptide (6 mg) dissolved in 3 mL of 50 mM sodium hydroxide solution was injected onto the SEC column. This normally elutes at a high concentration of approximately 250 µM. From roughly 60 mg of synthesized peptide, we collected about 30 mL of monomeric fractions. For oligomer preparation, monomers were incubated at 37 °C for 4 h. These incubated monomers were then reinjected into the SEC column, and the oligomer peak eluting at around 8 mL was collected. Oligomer fractions were kept on ice while the remaining monomer sample was processed. In total, approximately 12 mL of oligomer-containing fractions were obtained. This volume was concentrated, with approximately 70% of the buffer removed, using a centrifugal filter unit with a 3.5 kDa molecular weight cutoff (Amicon Ultra, Millipore) by centrifugation at 4 °C and 7500 rcf. The concentrated oligomers were then flash frozen in liquid nitrogen and lyophilized. The resulting material formed a fine white powder that was utilized for solid-state NMR experiments. This powder could also be readily resuspended in MilliQ water for downstream experiments.

#### SDS-PAGE and Silver Staining

To assess the molecular weight distribution of Aβ42 oligomer preparations, silver-stained SDS-PAGE was performed at room temperature (25 °C). Three lyophilized SEC fractions were reconstituted in phosphate-buffered saline (PBS): the SDS-induced 150 kDa oligomer fraction was dissolved in 50 µL PBS, the S1 fraction (an independently prepared batch of temperature-induced 37 °C oligomers) was dissolved in 20 µL PBS, and the S2 fraction (a second independent batch of 37 °C oligomers) was dissolved in 50 µL PBS. For each sample, 1 µL and 2 µL aliquots were loaded onto the gel without prior concentration determination. A Bio-Rad Precision Plus Protein™ standard was loaded as the molecular weight marker, and empty lanes were included as controls between sample groups.

Samples were resolved on a precast polyacrylamide gel under denaturing conditions. Following electrophoresis, gels were silver stained for high-sensitivity protein detection. The gel was fixed in a solution of 50% methanol and 5% acetic acid for 20 min, washed with 50% methanol for 10 min followed by water for 10 min, then sensitized in 0.02% sodium thiosulfate for 1 min. After rinsing with water, the gel was incubated in 0.1% silver nitrate solution for 20 min at 4 °C. Following additional water rinses, the gel was developed in 2% sodium carbonate containing 0.04% formaldehyde until bands reached the desired intensity, at which point the reaction was terminated with 5% acetic acid. Stained gels were imaged using a flatbed scanner or gel documentation system.

#### Cryo-Electron Microscopy

Negative staining and TEM imaging were performed following an approach similar to Kamalaldinezabadi et al.^18^. Following negative-stain screening, cryo-EM grids were prepared to further assess particle morphology and distribution at higher resolution. Lyophilized oligomer samples were resuspended in 10 mM sodium phosphate buffer (pH 7.5) immediately prior to grid preparation. Graphene-coated gold (Au) grids (UltraAuFoil R 1.2/1.3 Au300, Cat. #254401) were prepared by transferring a monolayer of graphene onto gold grids following a protocol established at the Stagg lab at Florida State University, available at https://www.secm4.org/faq/categories/protocols^53^. Grids were plasma cleaned using a Gatan Solarus (Model 950 Advanced Plasma System) prior to sample application. A 4 µL aliquot of sample was applied, followed by blotting and vitrification by plunge-freezing into liquid ethane using a Vitrobot at the FSU Biological Science Imaging Resource (BSIR) facility. Blotting conditions were set to a blot force of 1, blot time of 1 s, and single-sided blotting. A small cryo-EM dataset was collected on a Titan Krios transmission electron microscope (Thermo Fisher Scientific) operated at 300 keV and equipped with an Apollo direct electron detector at the FSU BSIR facility. Micrographs were acquired at a nominal magnification of 59,000×, corresponding to a calibrated pixel size of 0.79 Å/pixel, with an electron dose of ~60 e⁻/Å² and a defocus value of −3.0 µm.

#### Solid-State NMR Spectroscopy

Solid-state NMR experiments were performed on a Bruker AVANCE III HD-500 solid-state NMR system (500.13 MHz ¹H frequency) controlled with TopSpin 3.6.2 and equipped with an E-free 3.2 mm HCN triple-resonance magic-angle spinning (MAS) probe. Two-dimensional experiments were performed on 37 °C oligomers prepared with uniform ¹³C,¹⁵N labeling at L17, F20, A30, M35, V36 and I41. During the 2D exchange mixing periods, ¹³C–¹³C dipolar couplings averaged by MAS were reintroduced with the DARR sequence, using mixing times of 20 ms and 500 ms, at a sample-rotation rate of 10.5 kHz and with continuous-wave ¹H irradiation during mixing. 2D CHHC spectra were recorded at 10 kHz MAS with 150 µs ¹H–¹³C cross-polarization contacts before and after a 200 µs ¹H longitudinal mixing period, and 1D ¹H–¹³C CPMAS spectra were recorded at 10 kHz MAS. The sample temperature was set and maintained at room temperature (296–298 K) with Bruker temperature-control hardware, and spectrometer calibrations were performed using adamantane and/or polycrystalline glycine. ¹H–¹³C cross-polarization used a 2 ms contact time with a 50–100% ramp on the ¹H channel, and ¹H decoupling used two-pulse phase modulation (TPPM; 15° phase, 4.6 µs pulses) with a ¹H radio-frequency field of about 100 kHz during acquisition. Recycle delays were 1.5 s (DARR), 2 s (CHHC) and 4 s (CPMAS). The 2D DARR spectra were recorded as 2048 × 500 data points (direct × indirect) with spectral widths of 100 kHz and 50 kHz, and the CHHC spectrum as 2048 × 512 points with an indirect spectral width of 71.4 kHz. The 20 ms DARR spectrum is the result of 20.6 hours of signal averaging (96 scans per increment). The 500 ms DARR spectrum is the sum of three experiments of 27 hours each (96 scans per increment each). The CHHC spectrum is the sum of three experiments of 9.2 hours each (32 scans per increment each), each scaled by its stored intensity factor before summation. The 1D CPMAS spectra are the result of 7 minutes (labeled sample, 100 scans) and of 11.4 and 13.7 hours (the two unlabeled preparations, 10,240 and 12,288 scans) of signal averaging. The 2D spectra were processed in TopSpin with exponential line broadening of 50–100 Hz in the direct dimension (150 Hz in the indirect dimension of the CHHC spectrum) and a QSINC window (SSB 2) in the indirect dimension of the DARR spectra. Chemical shifts and linewidths in Table 1 were obtained from two-dimensional Gaussian fits to the 20 ms DARR spectrum.

### Supplementary Figures

#### Monomer Isolation from Synthetic Aβ42

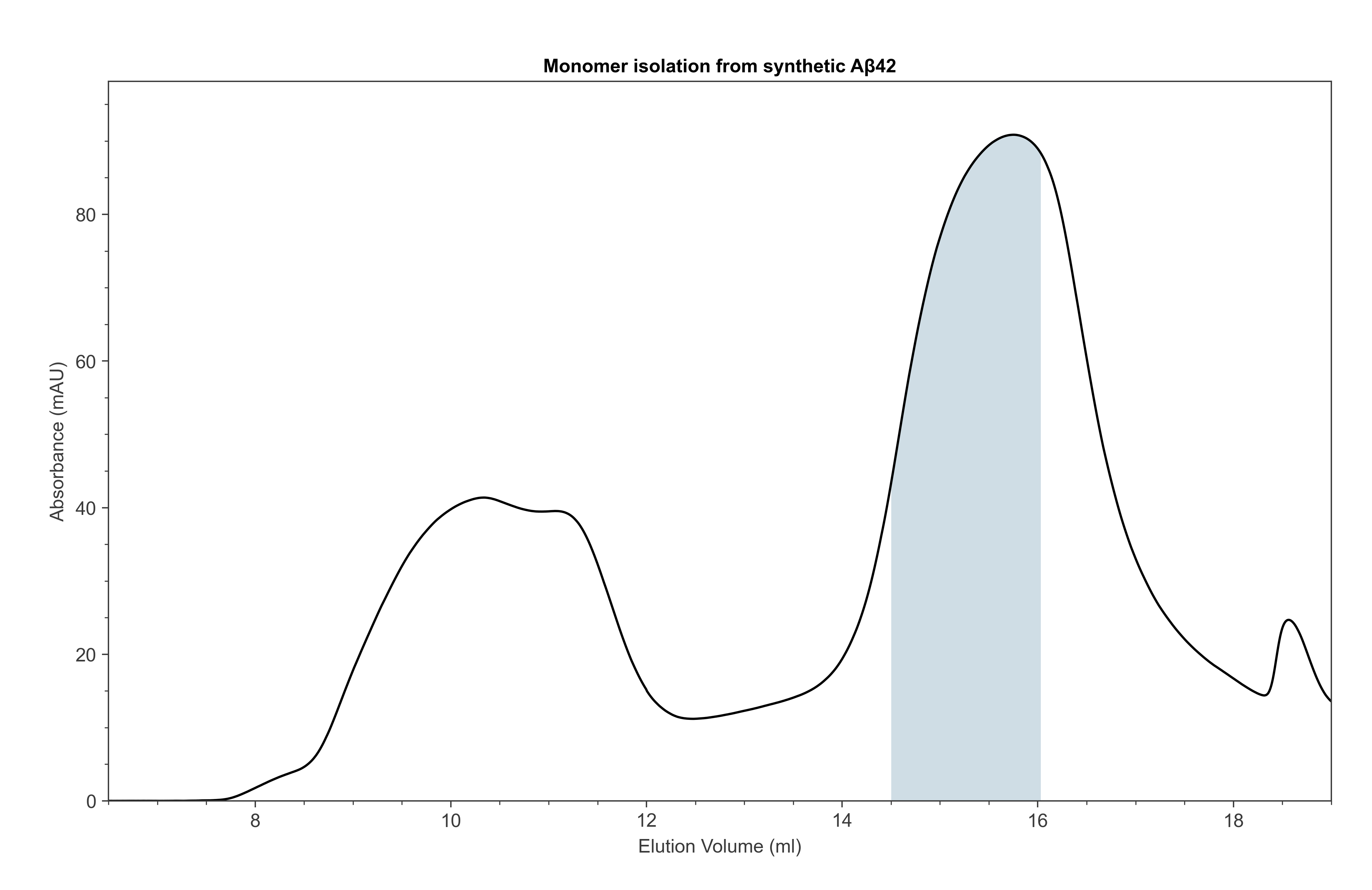

**Supplementary Figure 1. SEC profile measured for injection of synthetic Aβ42 prepared in 50 mM NaOH at a peptide concentration of 2 mg/mL, run on a Superdex 75 Increase column in 20 mM NaPi.** The shaded region (14.5–16.0 mL) marks the monomeric fractions collected for all subsequent experiments.

#### Thioflavin-T fluorescence of 25 µM monomers and oligomers

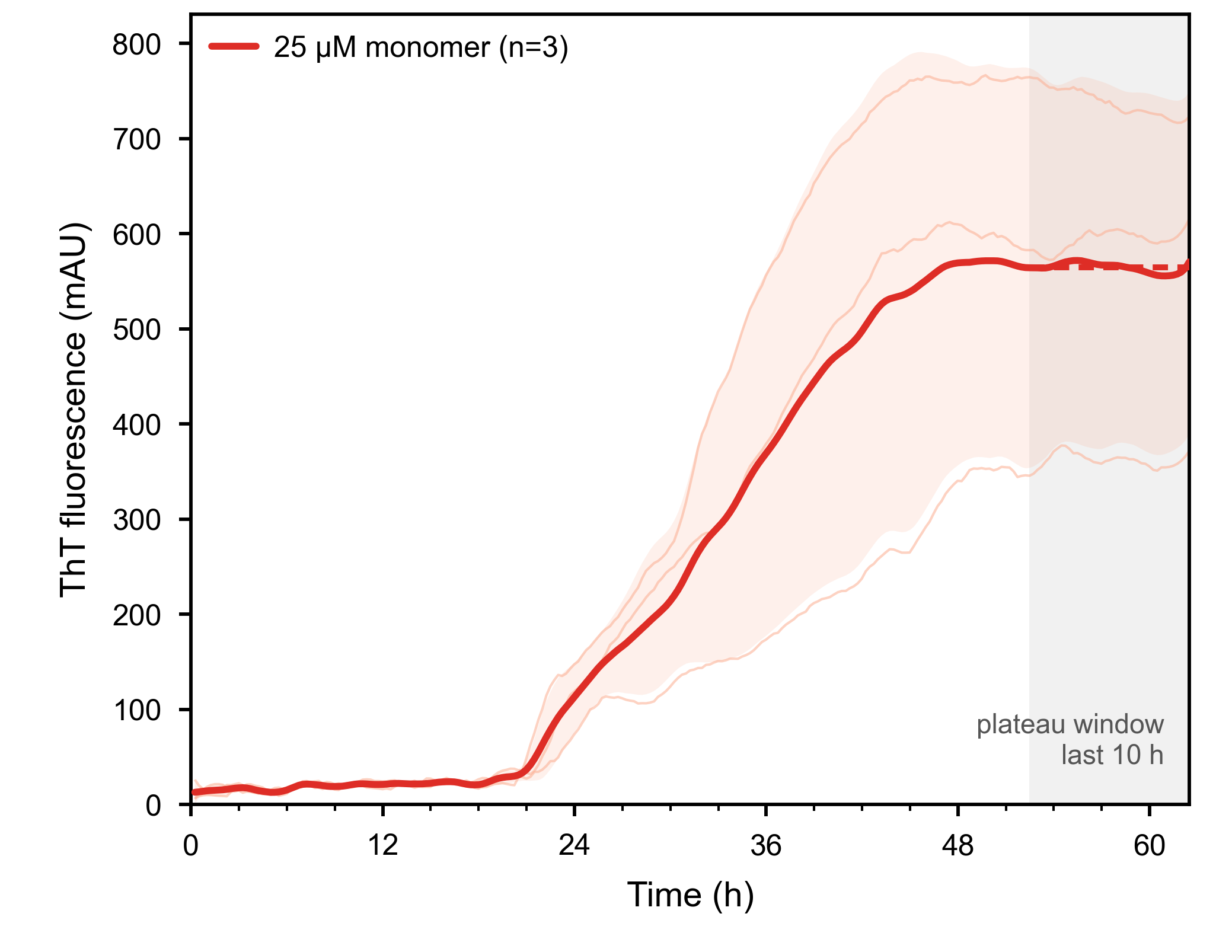

**Supplementary Figure 2. ThT fluorescence for 25 µM WT Aβ42 monomers at 37 °C with shaking, n = 3.** The solid line is the mean and the shading is ±1 SD. The gray band marks the plateau window, the last 10 h of the run, over which the fibril value in Fig. 1e is averaged.

#### Individual cytokine measurements in mouse microglial cells

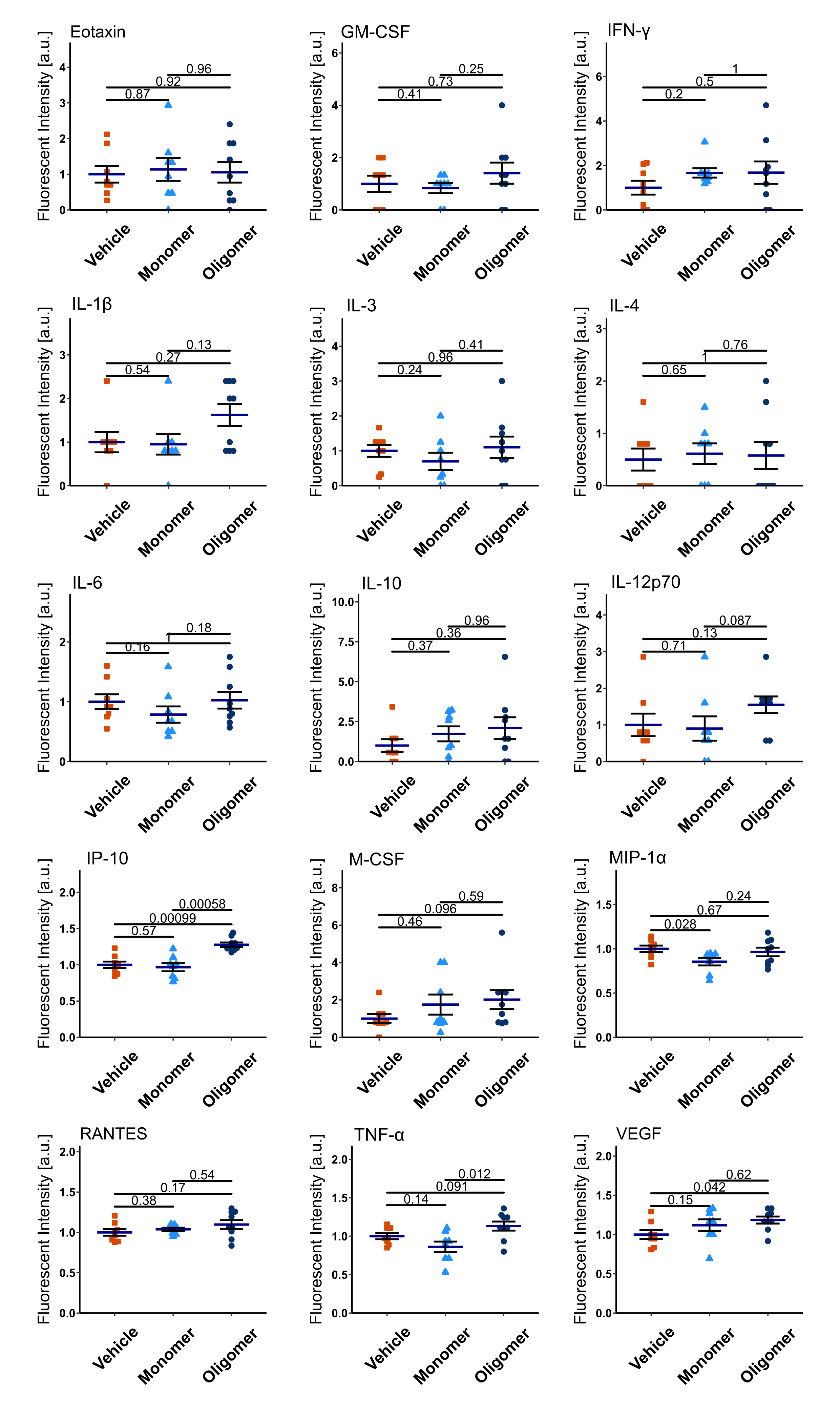

**Supplementary Figure 3. Individual cytokine responses of mouse microglial cells to Aβ42 monomers and oligomers.** Dot plots for all 15 cytokines (mean ± SEM) measured by multiplexed Luminex analysis of the MMC culture medium after 24 h of incubation with 50 nM of Aβ monomers, Aβ oligomers, or vehicle (n = 8–9 wells/group). TNF-α, VEGF and IP-10 are also shown in Fig. 2e. Each dot denotes an individual sample well. All p-values reflect Wilcoxon rank sum tests with Bonferroni adjustment for multiple comparisons. a.u., arbitrary units.

#### Individual cytokine measurements in hiPSC-derived microglia

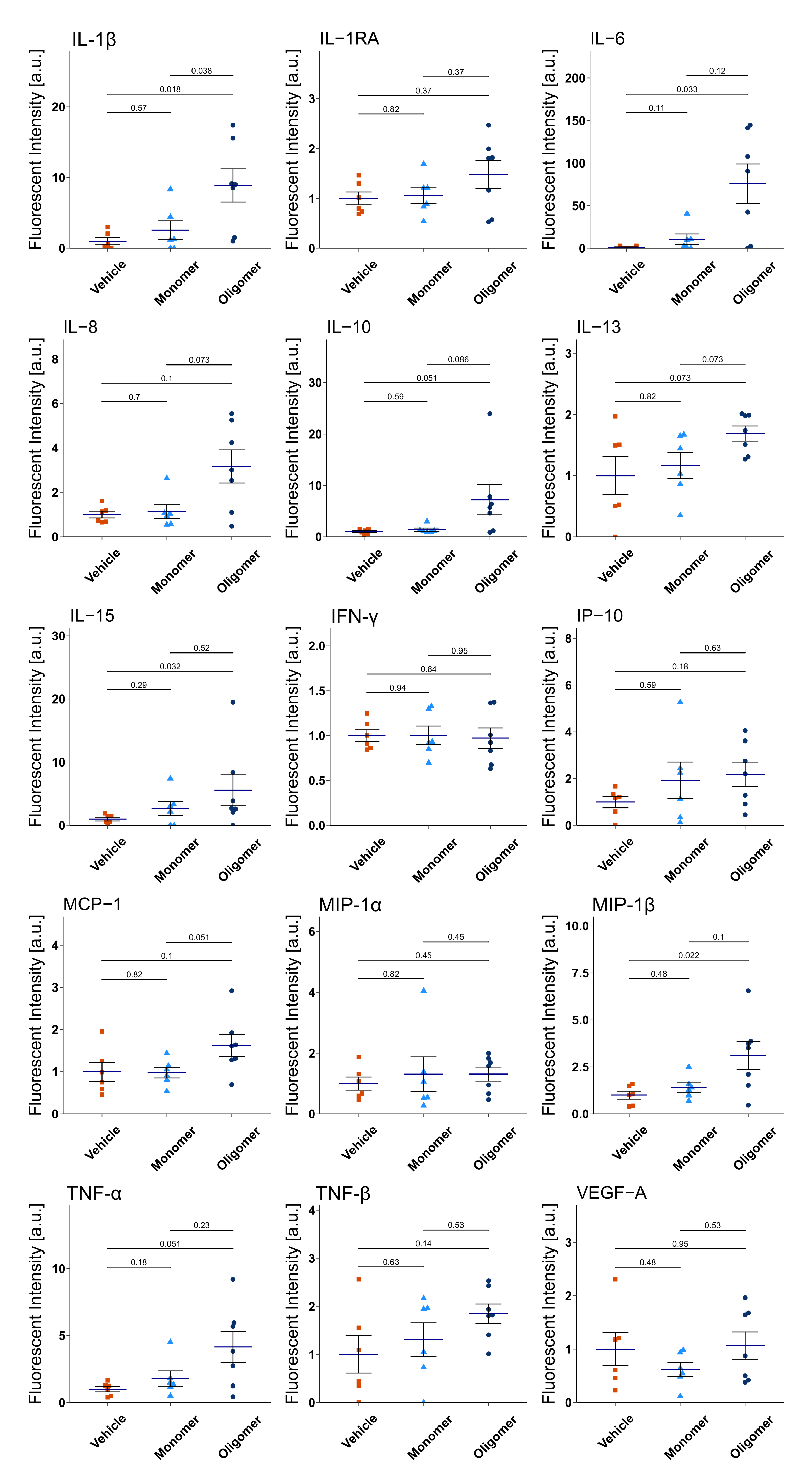

**Supplementary Figure 4. Individual cytokine responses of hiPSC-derived microglia to Aβ42 monomers and oligomers.** Dot plots for all 15 cytokines (mean ± SEM) measured by multiplexed Luminex analysis of the iMG culture medium after 24 h of incubation with 1 µM of Aβ monomers, Aβ oligomers, or vehicle (n = 6–7 wells/group). IL-1β, IL-6 and MIP-1β are also shown in Fig. 2j. Each dot denotes an individual sample well. All p-values reflect Wilcoxon rank sum tests with Bonferroni adjustment for multiple comparisons. a.u., arbitrary units.

#### Lower concentration of monomers impacts oligomer formation: Reanalysis at 210 nm UV detection wavelength

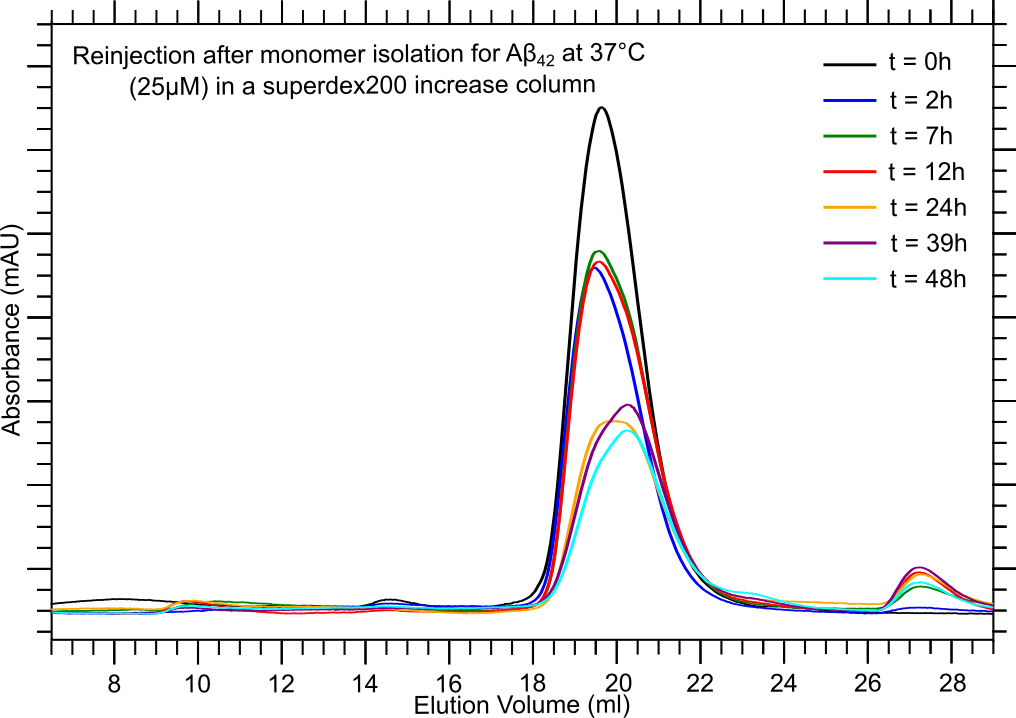

**Supplementary Figure 5. SEC profiles of 37 °C time-dependent monomer re-injections at a starting concentration of 25 µM, re-analyzed on a Superdex 200 Increase column with UV detection at 210 nm.** The peak at approximately 21 mL corresponds to monomeric Aβ42.

SEC profiles in Fig. 3 (main manuscript) demonstrate that Aβ42 oligomer formation depends nonlinearly on both concentration and temperature. When monomers were incubated at 37 °C for various time points at a low starting concentration of 25 µM, we observed little to no oligomer formation. The initial analysis was performed using a Superdex 75 Increase column with UV detection at 280 nm (Fig. 3c), and the absence of early-eluting peaks suggested that low monomer concentration reduces oligomer-forming propensity. To confirm this, we repeated the experiment using a Superdex 200 Increase column with UV detection at a more sensitive wavelength of 210 nm. The result showed the same pattern, with minimal peak intensity below ~16 mL elution volume, again indicating little to no oligomer formation at 25 µM.

#### Quantifying monomer and oligomer populations using SEC peak area

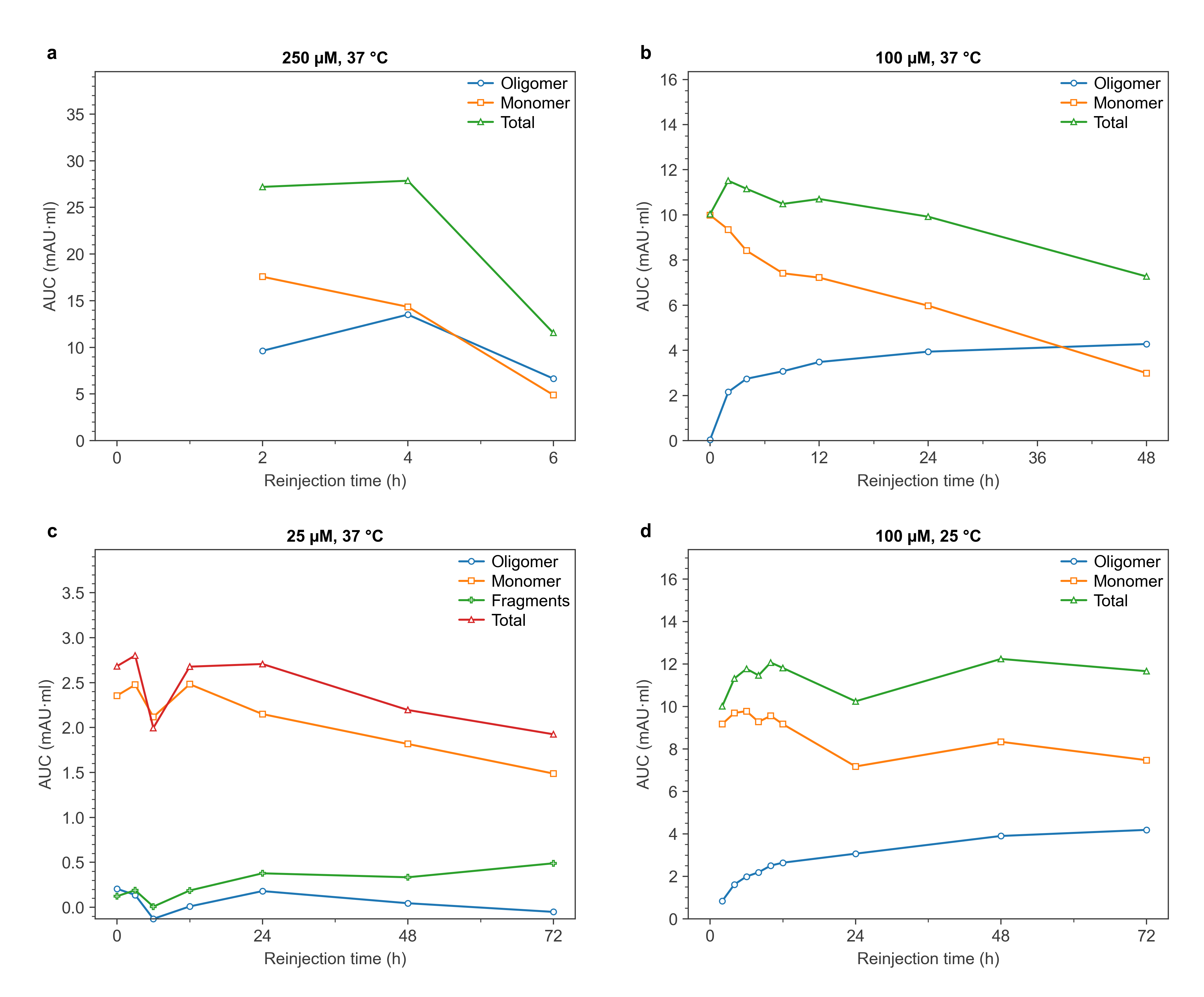

**Supplementary Figure 6. SEC peak area as a function of re-injection time for (a) 250 µM, 37 °C, (b) 100 µM, 37 °C, (c) 25 µM, 37 °C and (d) 100 µM, 25 °C.** Areas were obtained by trapezoidal integration of the raw absorbance, without baseline correction, over 5.0–11.0 mL (oligomer) and 12.0–18.0 mL (monomer). At 25 µM the monomer window is 12.0–16.1 mL and a separate 16.1–20.0 mL window captures the fragment peak near 17 mL. Total is the sum of the plotted components. Because no baseline is subtracted, areas for peaks near the noise floor can be slightly negative.

#### Repeatability of the oligomer preparation protocol

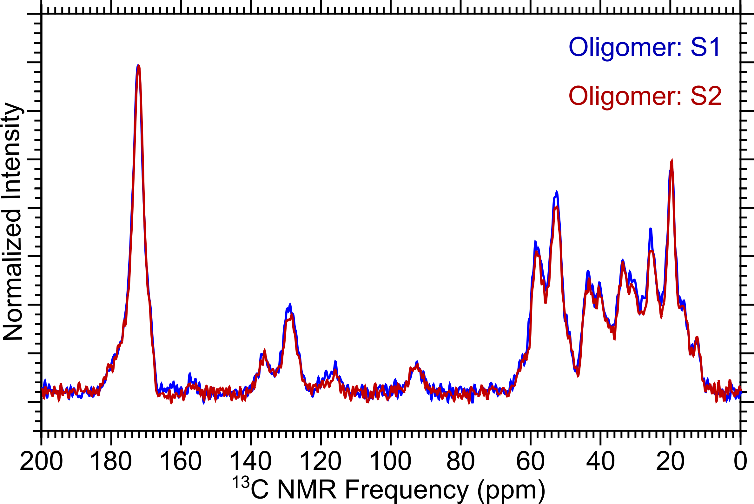

**Supplementary Figure 7. ¹H–¹³C CPMAS NMR spectra for two oligomer samples (S1 and S2) produced by the described protocol overlap, showing repeatability of the process.**

#### Solid-state NMR spectra

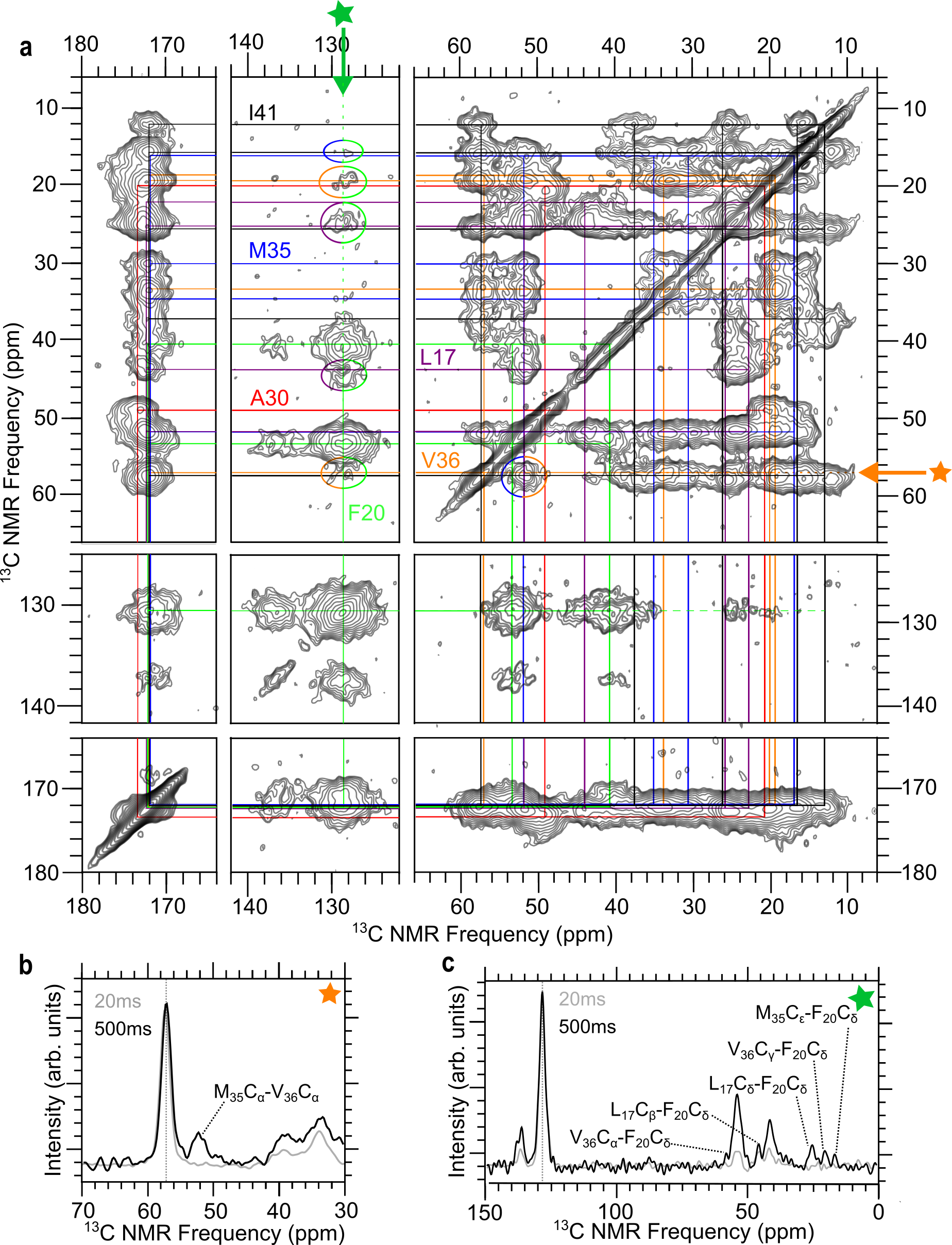

**Supplementary Figure 8. 2D ¹³C–¹³C DARR spectrum and slices for the 37 °C oligomer.** (**a**) Full 2D ¹³C–¹³C DARR spectrum at 500 ms mixing time, showing the aliphatic, aromatic and carbonyl regions with assignments for uniformly labeled L17, F20, A30, M35, V36 and I41. (**b**) Slice taken at 57.4 ppm, the V36 Cα frequency, showing the M35 Cα–V36 Cα crosspeak. (**c**) Slice taken at the F20 Cδ frequency, showing L17 Cβ–F20 Cδ, L17 Cδ–F20 Cδ, V36 Cα–F20 Cδ, V36 Cγ–F20 Cδ and M35 Cε–F20 Cδ. In both slices the 500 ms spectrum is black and the 20 ms spectrum gray.

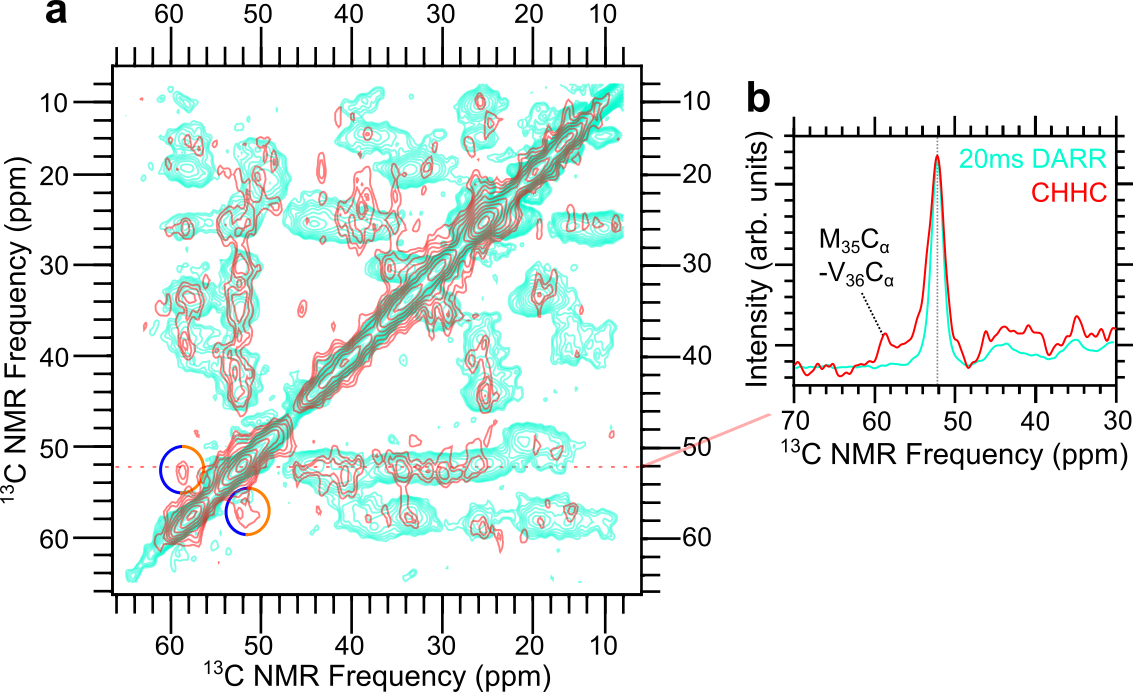

**Supplementary Figure 9. 2D CHHC spectrum of the 37 °C oligomer.** (**a**) Overlay of the 2D CHHC spectrum (red) on the 20 ms 2D DARR spectrum (cyan). Circles mark the M35 Cα–V36 Cα proximity on both sides of the diagonal. (**b**) Slice taken at 52 ppm showing a peak at 57.4 ppm present in the CHHC spectrum and absent from the 20 ms DARR spectrum.
